# Discovery and Predictive Modeling of Urine Microbiome, Metabolite and Cytokine Biomarkers in Hospitalized Patients with Community Acquired Pneumonia

**DOI:** 10.1101/2020.03.05.979427

**Authors:** Joseph F. Pierre, Oguz Akbilgic, Heather Smallwood, Xueyuan Cao, Elizabeth A. Fitzpatrick, Senen Pena, Stephen P. Furmanek, Julio A. Ramirez, Colleen B Jonsson

**Affiliations:** Department of Pediatrics, College of Medicine, University of Tennessee Health Science Center (UTHSC), Memphis, TN; Department of Health Informatics and Data Sciences, Parkinson School of Health Informatics and Public health, Loyola University Chicago, Maywood, IL, 60153; Department of Microbiology, Immunology, & Biochemistry, College of Medicine, UTHSC; Division of Infectious Diseases, School of Medicine, University of Louisville; Department of Acute and Tertiary Care, College of Nursing, UTHSC

**Keywords:** Microbiome, Metabolome, Urine, Predictive Modeling, Community-Acquired Pneumonia, Influenza, S aureus, S pneumoniae

## Abstract

Pneumonia is the leading cause of infectious related death costing 12 billion dollars annually in the United States alone. Despite improvements in clinical care, total mortality remains around 4%, with inpatient mortality reaching 5-10%. For unknown reasons, mortality risk remains high even after hospital discharge and there is a need to identify those patients most at risk. Also of importance, clinical symptoms alone do not distinguish viral from bacterial infection which may delay appropriate treatment and may contribute to short-term and long-term mortality. Biomarkers have the potential to provide point of care diagnosis, identify high-risk patients, and increase our understanding of the biology of disease. However, there have been mixed results on the diagnostic performance of many of the analytes tested to date. Urine represents a largely untapped source for biomarker discovery and is highly accessible. To test this hypothesis, we collected urine from hospitalized patients with community-acquired pneumonia (CAP) and performed a comprehensive screen for urinary tract microbiota signatures, metabolite, and cytokine profiles. CAP patients were diagnosed with influenza or bacterial (*S. aureus* and *S. pneumoniae*) etiologies and compared with healthy volunteers. Microbiome signatures showed marked shifts in taxonomic levels in patients with bacterial etiology versus influenza and CAP versus normal. Predictive modeling of 291 microbial and metabolite values achieved a +90% accuracy with LASSO in predicting specific pneumonia etiology. This study demonstrates that urine from patients hospitalized with pneumonia may serve as a reliable and accessible sample to evaluate biomarkers that may diagnose etiology and predict clinical outcomes.

**Author Summary:** Urine has been classically considered sterile since most microorganisms are not readily culturable under healthy circumstances. Further, many pneumonia patients are immediately placed on antibiotics rendering culture-based techniques useless. However, the advent of next generation sequencing has enabled unprecedented analysis of the microbial communities – living or detected as free DNA – found in many niches of the human body. Here, we describe a urine microbiome as well as metabolites and cytokines measured in patients newly admitted to the hospital diagnosed with influenza or bacterial (*S. aureus* and *S. pneumoniae*) infection pneumonia, compared with healthy controls. Using these parameters alone, we were able to achieve high success in predicting patient pneumonia. This study provides a proof of concept that urine samples, which are easily accessible in outpatient and inpatient settings, could provide additional diagnostic insights to patient infectious status and future risk factor for complication.

## Introduction

Community-acquired pneumonia (CAP) is the leading cause of infectious disease-related death and together with Influenza, the eight-leading cause of death in the USA ^1^. The annual incidence of CAP worldwide is approximately 5-11 per 1000 and the estimated annual CAP-associated costs in the US is over 12 billion dollars ^2^. Even with appropriate antibiotic and Supportive therapy, some hospitalized patients with CAP progress to clinical failure and death ^3^. Intriguingly, even patients that survive the initial respiratory infection have significantly higher 1, 3, and 5-year mortality rates compared to other chronic diseases (reviewed in ^4^). Therefore, there is a critical need to improve treatment and gain a deeper understanding into the factors contributing to short and long-term morbidity and mortality.

Both bacterial and viral pathogens cause CAP and both etiologies are associated with significant mortality ^3, 5, 6^. Prompt identification of the pathogen causing pneumonia is critical for prescribing appropriate therapy. However, the tests necessary to identify the pathogen in blood or bronchoalveolar lavage (BAL) suffer from sensitivity, specificity, cost and availability issues ^5, 7–9^. Frequently, no pathogen is identified making treatment decisions exceedingly difficult and adversely affecting patients; delays in antibiotic treatment are associated with increased mortality. Additionally, the symptoms of viral and bacterial pneumonia overlap and it is difficult to distinguish between the two based on clinical and radiographic findings^10–13^. Early identification of the etiology is critical for prescribing appropriate treatment; unfortunately, there is no standard diagnostic criterion for distinguishing between viral vs bacterial pneumonia. There have been numerous attempts to identify biomarkers that will distinguish viral versus bacterial pneumonia however none have become part of standardized hospital diagnosis practice.

Many studies have quantified cytokines or eicosanoids in serum as potential biomarkers for pneumonia severity. According to the current paradigm, high cytokine levels produce an exaggerated systemic response – termed the “cytokine storm” - and this dysregulated systemic inflammation drives poor clinical outcomes ^14^. However, using cytokines alone as biomarkers for severity has not been well established or incorporated into standard diagnostic criteria. In addition to cytokines, the advent of next generation sequencing has allowed rapid identification of specific microbiome compositions in multiple body sites. While urine was classically considered sterile, recent reports suggest a unique microbiome is detectable under healthy and diseased conditions. ^15, 16^ These microbes, along with the mammalian host, produce a milieu of metabolites. Metabolites are functional outputs of various biological processes and they are an end point that incorporates biological state of the patient (e.g. age and genetic factors) with disease state and external environmental influences (e.g. nutrition and drug treatment) and internal microbiome influences. Metabolites are dynamic analytes present in biological fluids, including urine, producing unique signatures that are readily detected with mass spectrometry and are currently being studied in patients with pneumonia ^17–20^. While these studies have made great strides in elucidating underlying mechanisms of pneumonia and determined some promising biomarkers for specific causative agents, none have combined metabolomics and microbial-omics nor expanded from the discovery phase to algorithmic predictive modeling.

The metabolome and microbiome ^21^ are also unique portraits of the individual patient as they are not only influenced by genetics and disease state but also by the environment, nutrition, age, and lifestyle ^19, 22, 31, 23–30^. Thus the microbiome and metabolome are very different from transcriptomic or proteomic biomarkers. Unlike biomarkers that individually vary with health status, meta-biomarkers by definition are so co-related and interwoven that they produce a precise disease signature that evolves with the pathophysiological state of the individual ^31–34^. Thus meta-biomarkers are less likely to produce false identifications as they do not rely on changes to a single analyte ^31, 34^. Temporal and dynamic changes to the microbiome and metabolome along with their connection to phenotype and meta-biomarker characteristics leads us to select these quantifiable components in the urine as indicators of CAP. Therefore, here we set out to identify whether unique signatures of patients with CAP could be identified in urine by incorporating cytokines, the microbiome, and metabolites in our predictive models.

## Results

### Patient Characteristics

Patients were selected from the University of Louisville Pneumonia Study (ULPS) biorepository with IRB approval. The ULPS was a population-based cohort study of 7,449 unique patients hospitalized with CAP between June 1, 2014 and May 31, 2016. From this biorepository, we selected 30 urine samples from patients, ten each with a confirmed etiology of influenza A virus (IAV), *S. pneumoniae,* or *S. aureus* infection. **Table 1** shows a comparison of clinical data for patients hospitalized with CAP for each etiologic agent. We also selected urine samples from ten healthy volunteers from the University of Louisville Infectious Diseases biorepository. These volunteers were majority female (90%) and aged 28-58.

**Table 1.** Clinical characteristics of hospitalized patients due to CAP.

| Variable | Influenza | *S. aureus* | *S. pneumoniae* |
| --- | --- | --- | --- |
| n | 10 | 10 | 10 |
| **Demographics** |  |  |  |
| Age (median (IQR)) | 54 (50-62) | 61 (50-75) | 61 (44-63) |
| Male Sex (%) | 7 (70) | 5 (50) | 5 (50) |
| Caucasian (%) | 8 (80) | 10 (100) | 8 (80) |
| **Social and Medical History** |  |  |  |
| Neoplastic disease (%) | 2 (20) | 2 (20) | 1 (10) |
| Congestive heart failure (%) | 4 (40) | 1 (10) | 1 (10) |
| Stroke (%) | 0 (0) | 3 (30) | 0 (0) |
| Renal disease (%) | 4 (40) | 3 (30) | 0 (0) |
| Liver disease (%) | 0 (0) | 1 (10) | 1 (10) |
| Diabetes (%) | 3 (30) | 3 (30) | 3 (30) |
| COPD (%) | 5 (50) | 4 (40) | 4 (40) |
| Human immunodeficiency virus | 1 (10) | 0 (0) | 0 (0) |
| Active coronary artery disease (%) | 3 (30) | 1 (10) | 2 (20) |
| Arterial hypertension (%) | 8 (80) | 7 (70) | 7 (70) |
| Hyperlipidemia (%) | 6 (60) | 5 (50) | 1 (10) |
| Prior myocardial infarction (%) | 3 (30) | 0 (0) | 1 (10) |
| Prior PTCA/CABG (%) | 3 (30) | 1 (10) | 1 (10) |
| Atrial fibrillation (%) | 2 (20) | 1 (10) | 1 (10) |
| Obese (%) | 6 (60) | 0 (0) | 5 (50) |
| Smoking status (%) |  |  |  |
| Current | 5 (50) | 6 (60) | 5 (50) |
| Former | 5 (50) | 2 (20) | 2 (20) |
| Never | 0 (00) | 2 (20) | 3 (30) |
| **Physical Examination and Laboratory Findings** |  |  |  |
| Heart rate (Beats/Minute) | 118 (110-125) | 112 (103-115) | 113 (103-131) |
| Respiratory rate (Breaths/Minute) | 28 (26-30) | 27 (21-35) | 26 (21-33) |
| Systolic blood pressure (mmHg) | 122 (103-151) | 109 (88-123) | 102 (82-124) |
| Diastolic blood pressure (mmHg) | 58 (49-65) | 50 (43-58) | 54 (42-67) |
| Temperature (Degrees Celsius) | 38 (38-39) | 38 (38-39) | 38 (37-38) |
| Hematocrit(%) | 40 (39-41) | 28 (25-32) | 40 (31-40) |
| Serum sodium (mEq/L) | 135 (133-138) | 129 (126-134) | 135 (133-136) |
| Blood Urea Nitrogen (BUN) (mg/dL) | 19 (13-23) | 25 (18-40) | 15 (13-24) |
| Serum bicarbonate (mEq/L) | 26 (24-27) | 22 (19-26) | 26 (25-28) |
| Serum glucose (mg/dl) | 112 (91-156) | 149 (119-241) | 124 (117-150) |
| PaO2/FiO2 < 250 (%) | 4 (40) | 3 (30) | 4 (40) |
| Multilobar Pneumonia (%) | 6 (60) | 6 (60) | 5 (50) |
| Pleural Effusion (%) | 2 (20) | 4 (40) | 3 (30) |
| **Severity of Disease** |  |  |  |
| Pneumonia severity index class IV/V | 6 (60) | 7 (70) | 6 (60) |
| Non-invasive mechanical ventilation on day 0 (%) | 0 (0) | 1 (10) | 1 (10) |
| Invasive mechanical ventilation on day 0 (%) | 1 (10) | 2 (20) | 0 (0) |
| Vasopressors on day 0 (%) | 0 (0) | 1 (10) | 0 (0) |
| Altered mental status on day 0 (%) | 3 (30) | 1 (10) | 2 (20) |
| **Clinical Course** |  |  |  |
| Acute MI during hospitalization (%) | 0 (0) | 1 (10) | 0 (0) |
| Need for invasive ventilation after day 0 (%) | 0 (0) | 2 (20) | 1 (10) |
| Need for vasopressors after day 0 (%) | 1 (10) | 2 (20) | 2 (20) |
| Time to first antibiotics (hours) | 2.8 (2.3-3.8) | 2.1 (1.8-3.1) | 2.8 (2.0-3.5) |
| Time to Clinical Stability (days) | 2 (2-2) | 5 (2-7) | 2 (2-5) |
| Length of Stay (days) | 4 (3-4) | 8 (7-13) | 5 (4-9) |
| **Outcomes** |  |  |  |
| Rehospitalization due to CAP (%) | 1 (10) | 0 (0) | 0 (0) |
| Mortality during hospitalization (%) | 0 (0) | 0 (0) | 1 (10) |
| Mortality at 30 days (%) | 0 (0) | 1 (10) | 1 (10) |
| Mortality at 6 months (%) | 0 (0) | 1 (10) | 1 (10) |
| Mortality at 1 year(%) | 0 (0) | 1 (10) | 3 (30) |

### Urine Cytokines

We interrogated 34 cytokines in the urine of healthy controls, patients infected by influenza, *S. pneumo,* or *S. aureus*. Out of 34 cytokines tested, we detected 17 cytokines in the urine and 11 of those cytokines showed differential presentation among the four groups of participants (**Table 2**). Pneumonia caused by *S. aureus* differed from the healthy controls for all 11 cytokines and in general the level of cytokines detected in influenza patients was consistently lower than the levels in patients with bacterial pneumonia and only differed from healthy volunteers for IFNγ, IL-6, IL-18, eotaxin, IP-10 and MCP-1. Four of the cytokines demonstrated a significant difference between the 3 types of pathogens; IFNγ (P=0.005), IL-18 (P=0.0052), MCP-1 (P=0.0029) and SDF-1 (P=0.0451). The remaining cytokines that were present in the urine but did not differ from each other or healthy volunteers were IL-1β IL-1α IL-1RA, IL-22, IL-27 and IL-8. We did not detect IL-12p70, IL-13, IL-2, IL-5, GM-CSF, IL-10, IL-17A, IL-21, IL-23, IL-9, IFNα, IL-31, IL-7, TNFβ MIP-1, RANTES or TNFα in the urine in any of the groups.

**Table 2.**
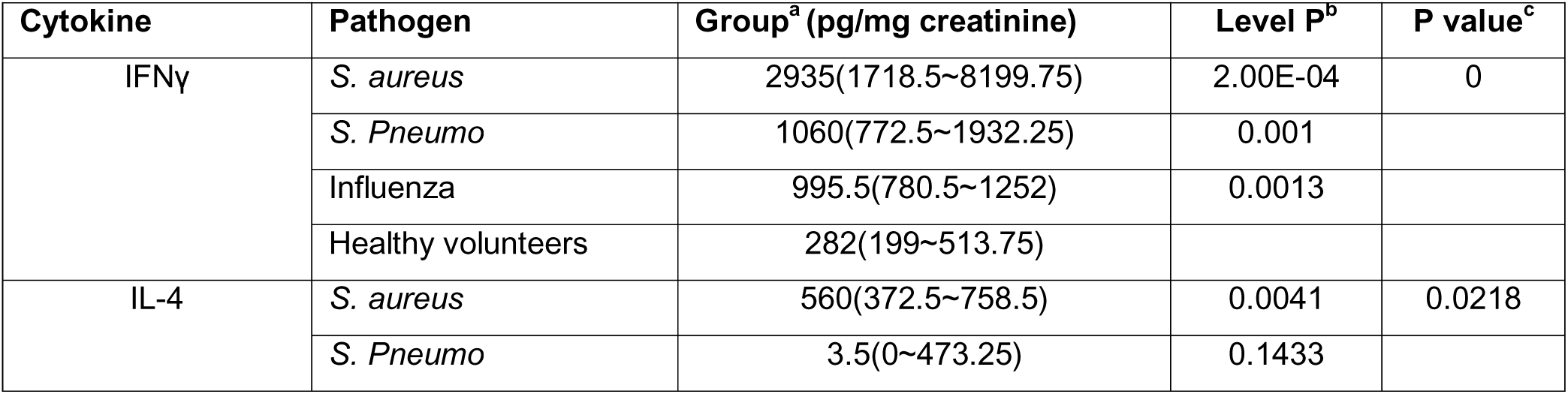
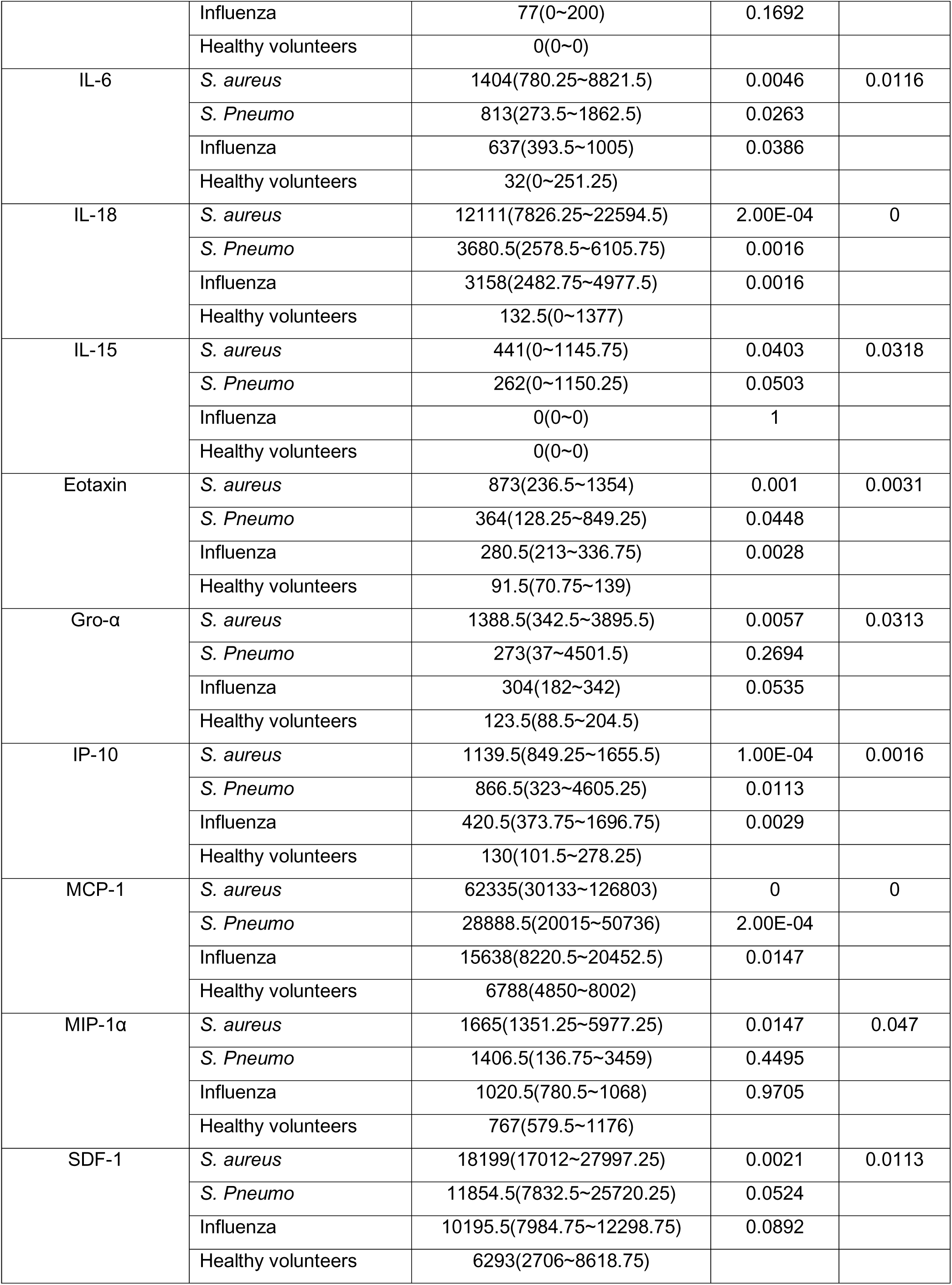
Cytokines detected in the urine of CAP patients

### Urine Microbiome

Total DNA was extracted from urine samples for quantitative PCR of 16S copy numbers, which demonstrated that healthy volunteers exhibited higher bacterial DNA copy numbers compared with bacterial or viral pneumonia patient urine samples (**Fig. 1A**); potentially influenced by the initial administration of antibiotics all cause pneumonia patients receive upon admission. Similarly, compared with healthy volunteers, pneumonia patients tended to have elevated alpha diversity, assessed by Shannon index (Anova: F=2.5, P=0.076) (**Fig. 1B**) and significantly elevated Evenness (Anova: F=3.3, P=0.03) (**Fig. 1C**). Microbiome taxonomic composition across individuals displayed taxonomic signatures of pneumonia, with elevated phyla Proteobacteria and Chloroflexi and fewer Firmicutes compared with healthy controls (**Fig. 1D, S1A Fig.**). Genera level taxa are shown in **Fig. 1E** across individuals. Principal component analysis (PCoA) of beta diversity assessed with Bray-Curtis broadly demonstrated clustering of pneumonia patients compared with healthy volunteers (**Fig. 2A**). Redundancy Analysis (RDA) further demonstrated similar distinct clustering between healthy and influenza samples, while patients infected with bacterial pathogens, *S aureus* and *S pneumonia*, clustered even more distinctly from healthy volunteer samples (RDA significance: Variance=12.74, F=1.13, P=0.05) (**Fig. 2B**). A network analysis of detected taxa also demonstrated more tightly clustering of taxa found in healthy volunteer associated (**Fig. 2C**). Specific taxa associated with healthy volunteers included the genus *lactobacillus* within Firmicutes, while pneumonia patients demonstrated generally elevated levels of the class *Gammaproteobacteria* and specifically *Clostridium* and *Enterobacteriaceae* (**Fig. 2D, S1 B Fig. 1B, S2A,B Fig.**). Hierarchical taxonomic composition for each patient group are displayed in **S3 Fig**.

**Figure 1.**
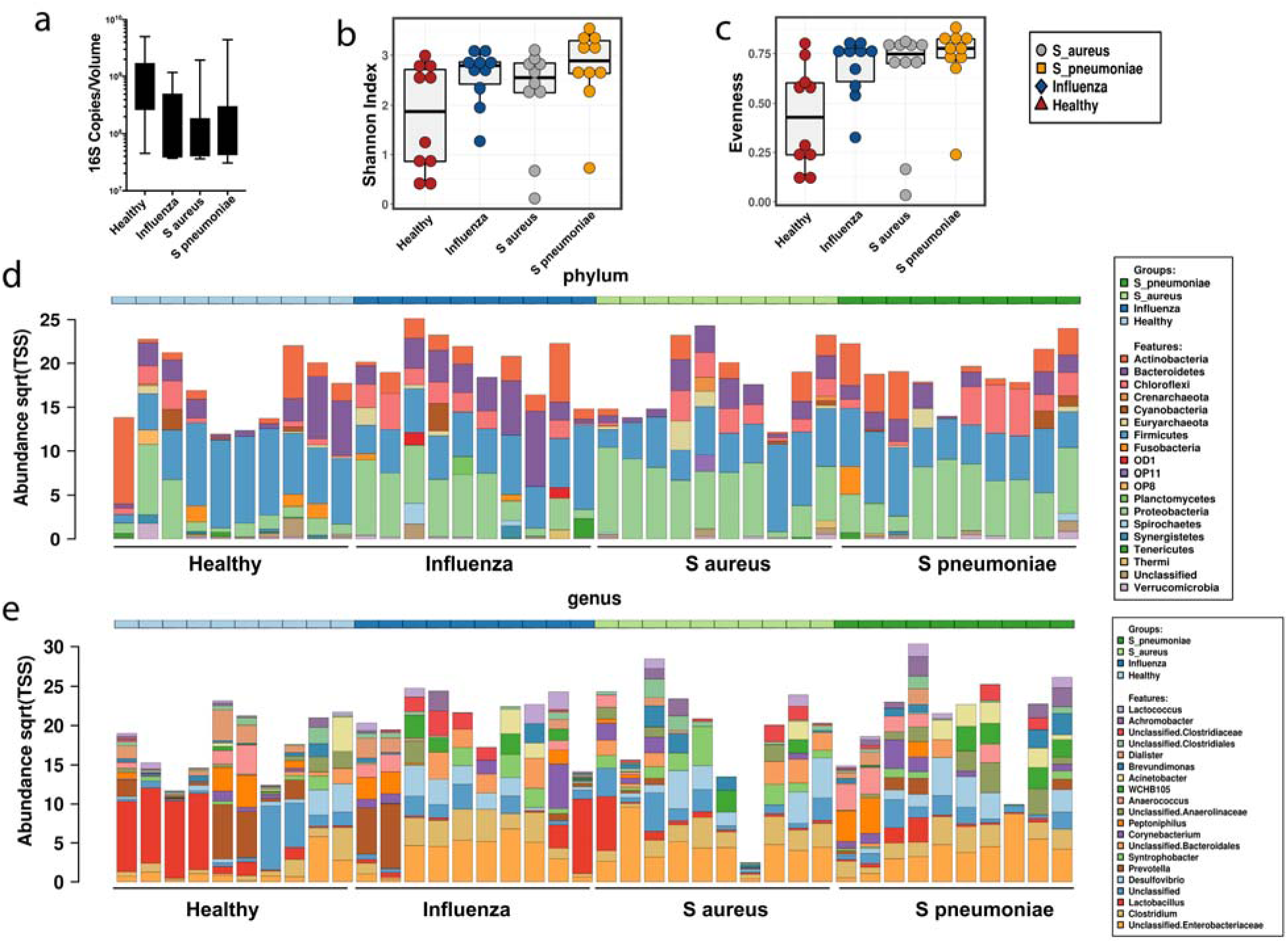
Urine Microbiome Alpha Diversity of Taxonomic Analysis. (a) 16S copy numbers detected per mL of urine. (b) Shannon and (c) evenness assessment of alpha diversity. Taxonomic community structure of each sample at phylum (d) and genus (e) level. N=10/group.

**Figure 2.**
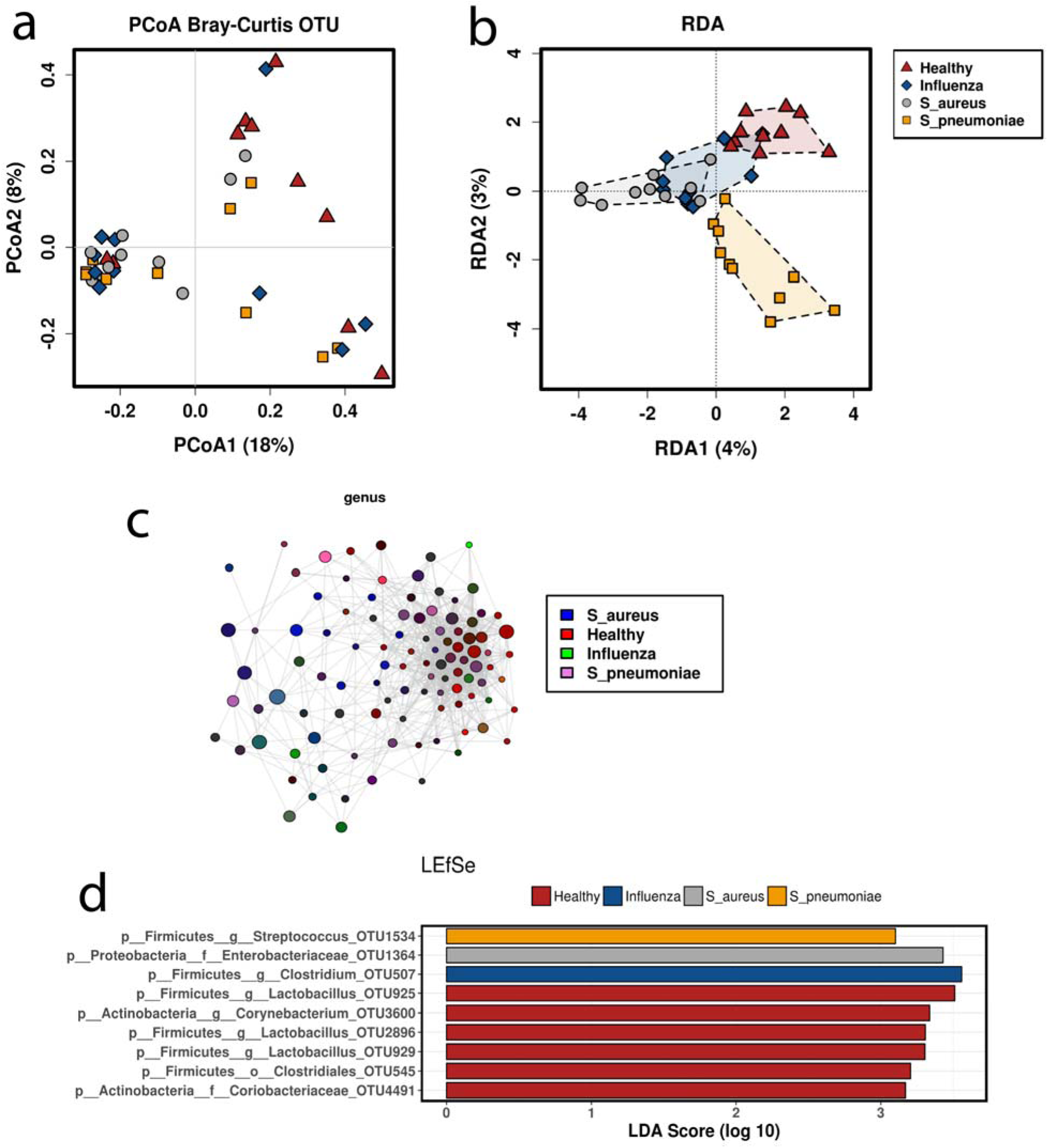
Urine Microbiome Beta Diversity, Network clustering, and LEfSe. (a) Principal component analysis of Bray-Curtis beta diversity of urine OTUs. (b) Redundancy analysis of urine OTUs. (c) Network analysis of genus detected in urine, color coded by group. (d) Linear Discriminant Analysis of Effect Size (LEfSe) of OTUs enriched in each experimental group. N=10/group.

To identify specific taxanomic differences between groups, we further employed linear discriminate analysis of effect size (LEfSe) between experimental groups. Initially, we compared all 30 patients with CAP to healthy volunteers (**S4A Fig.**). At the phylum level, Proteobacteria was identified as significant in pneumonia samples while *Synergistetes* was identified in healthy controls based on LDA scores. At the genus level, *Clostridium* and *Sutterella* were identified in case with CAP while *Lactobacillus, Prevotella, Magasphaera, Dorea, Vibrio*, and *Coprococcus* were the most significantly enriched. Cladogram projection of these differences demonstrated the CAP case samples clustered primarily within the phyla Proteobacteria, while the healthy volunteers were more taxonomically distributed (**S4B Fig.**). The analysis was repeated after regrouping samples based on viral vs bacterial pathogen, which showed changes at the order level, where *Bifidobacteriales* was abundant in healthy controls, *Enterobacteriales* was abundant in bacterial pneumonia samples, and *Sphingomonadales* was abundant in influenza samples (**S4C Fig.)**. A final regrouping determined comparisons of healthy vs *S. aureus* and *S. pneumoniae* (**S4D Fig.**) and healthy vs influenza (**S4E Fig.**), where healthy samples consistently displayed greater levels of the family Rikenneliaceae and the order Bifidobacterium. Interestingly, the level of detectable *Streptococcus* was most elevated in *S pneumonia* case samples, while *Syntrophobacter* and *Delftia* were most elevated in patients with *S aureus* (**S2B Fig.**).

### Urine Metabolites

Metabolites were extracted from 50 µl of urine and subjected to ultra-high-performance liquid chromatography coupled high resolution mass spectrometry (UPLC-HRMS) 87 known metabolites were detected in the forty urine samples and identified using known masses (+/- 5 ppm) and retention times (Δ ≤1.5 min). Creatinine is considered the best internal standard to correct for urine volume variations as its rate of elimination is independent of urine flow and urine volume and creatinine concentration are inversely proportional ^35, 36^. Thus, to ensure that observations were directly comparable peak intensity was normalized to creatinine. Then these data were compared to unnormalized data to make sure there was no masking of biologically relevant changes by normalization (DNS). As is the convention in metabolomics we first used unsupervised multivariate statistical analysis to determine the dataset structure and relationships between groups.

To evaluate the group trends, sample uniformity and identify potential outliers, multivariant principal component analysis (PCA). The variation were explained by F1 and F2 with a cumulative percent variability of 78.56% spread among the patient groups (i.e. Healthy 53.5 and 6.1, IAV 13.8 and 70.0, *S. aureus* 8.6 and 22.4, and *S. pnuemo* 30.0 and 1.2 percent per F1 and F2 respectively). Adding a third component marginally increased the cumulative percent variability to 85.96%. The two component PCA analysis shows good separation between CAP patients and the healthy group (**Fig. 3, circles)**. Likewise, the high-risk classes (IV-V) and low risk (I-III) centroids showed clear separation (**Fig. 3, squares)**. We then used unsupervised clustering of both metabolites and individuals, they were clustered independently using k-means clustering followed by ascendant hierarchical clustering based on Euclidian distances. The data matrix’s was rearranged according to the corresponding clustering with spatial relationship proportional to similarity among patient samples or metabolites **(Fig. 4)**. These clusters were also represented via a dendrogram displayed vertically for metabolites and another horizontally for patients. We find the healthy volunteers centered and groups nicely together (red) as did the IAV (Blue) while the bacterial pneumonia samples were interspersed together (gray and orange) (**Fig. 4**). Consistent with the PCA analysis the high-risk groups tended to be close together on the far left or right (**Fig. 4 brown bars**).

**Figure 3.**
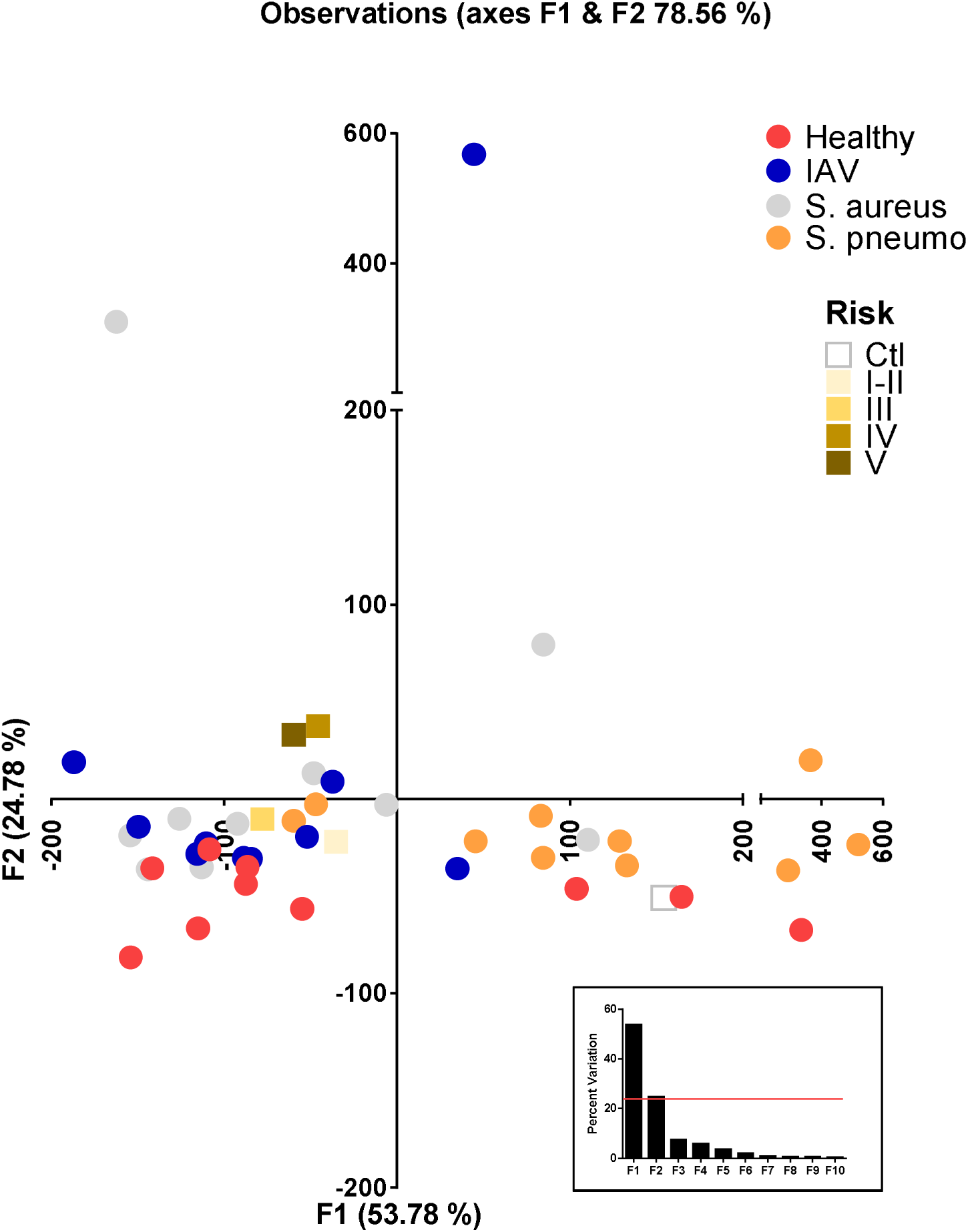
Principal component analysis of urine metabolites. Metabolites were extracted from 50µl urine and subjected to UPLC–HRMS metabolomics analysis three times per sample. Metabolites were manually identified and integrated using known masses (± 5 ppm mass tolerance) and retention times (Δ ≤ 1.5 min). Peak intensity was normalized to creatinine followed by unsupervised multivariant principal component analysis (PCA) resulting in F1 and F2 with a cumulative percent variability of 78.56% Each circle represents the average of a patient and the centroids of the corresponding risk groups are represented by squares.

**Figure 4.**
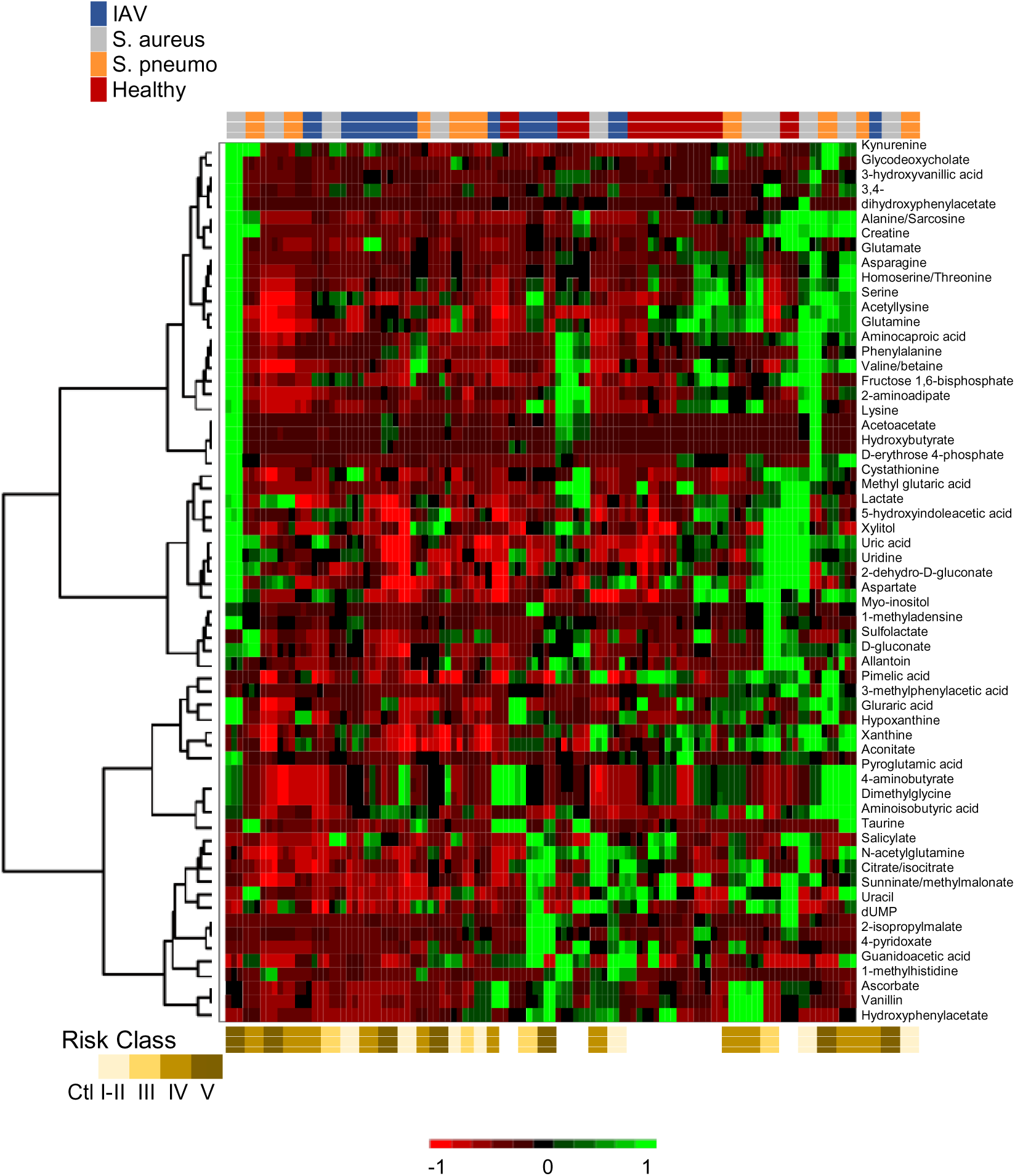
Comparison of urine metabolites by patient and risk group. Metabolites were K-means clustered followed by ascendant hierarchical clustering based on Euclidian distances with twenty-one metabolites excluded (0.25 < std dev). Metabolite clusters were also represented via a dendrogram displayed vertically for metabolites and another horizontally for patients. The data values of the permuted matrix were replaced by corresponding color intensities based on interquartile range with color scale of red to green through black resulting in a heat map. Patient identifiers and risk categories were replaced by color bars. Color bars on the top of the graph denote patient groups and bottom risk class.

Next, we employed a simple one-way ANOVA with Tukey’s honestly significant difference test (Tukey’s HSD) with Benjamini-Hochberg post hoc correction (XLSTAT) to identify 6 metabolites with significant differences among patient groups (**Table 3**). Adenosine 5‘-phosphosulfate (APS) was the most significant metabolite with differential concentration based on pneumonia, it was significantly higher in the urine of healthy volunteers. In humans, all APS is converted to 3’-phosphoadenosine 5’-phosphosulfate (PAPS) for the sulfonation of glycosaminoglycans, proteins, peptides, lipids, bile acids, xenobiotics and steroids ^37–39^. Guanidoacetic acid was also significantly higher in healthy volunteers and is a precursor to creatine, metabolite in the Urea cycle as well as metabolism of amino groups of several amino acids including glycine, serine, threonine, arginine and proline. 2,3-dihydroxybenzoate is a conjugate base of 2,3-dihydroxybenzoic acid that is increased after consumption of nutrients (e.g. cranberry juice) or aspirin and is also a biomarker of OH radicals ^40, 41^. Succinate was significantly decreased in patients with pneumonia in our studies as well as two other metabolite profiling studies of pneumonia from human pleural fluid and mouse urine infected with *S. pneumoniae* ^18, 42^. We found citrate and succinate, metabolites related to the citric acid cycle, to be significantly reduced in all three groups with CAP (**Table 3**). Reduced citrate levels have previously been reported in plasma of patients with pneumonia and in mouse urine ^18, 27^. Likewise, Adamko and co-workers observed decreases in both citrate and succinate in urine from children with bacterial and viral respiratory infections ^18, 27, 43^. Conversely, we found uridine to be significantly increased in the urine of all patients with pneumonia. This is in keeping with previous reports of uridine transiently increasing in the lung and BAL fluid of mice with viral pneumonia from influenza ^42^. It is worth noting these are highly abundant analytes whose values are relative to the peak intensity of creatinine in each sample (creatinine mean across samples was 2.753e+009). Taken together these metabolites represent likely candidates for including among the signature biomarkers of pneumonia.

**Table 3.**
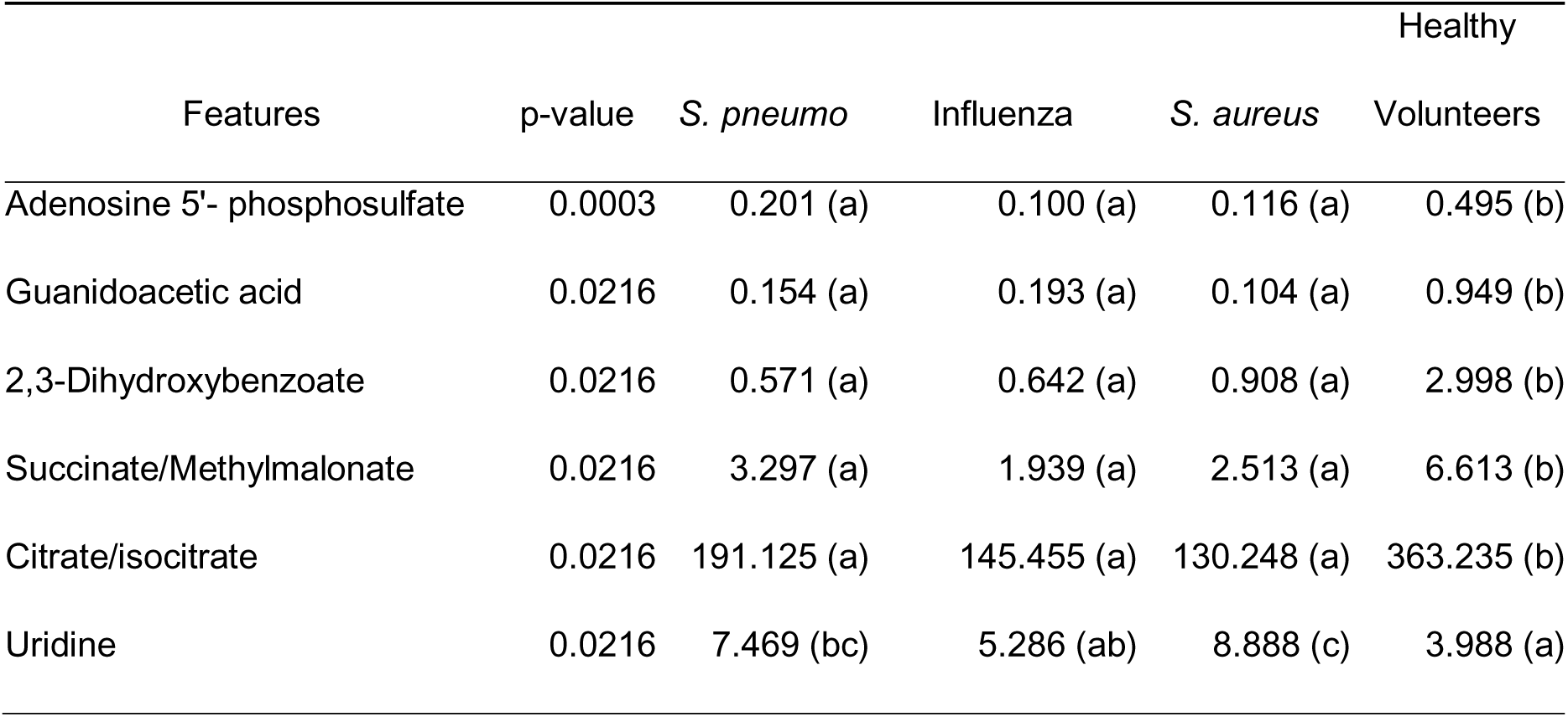
Metabolites differentially observed between groups

We applied a supervised four component partial least squares discriminant analysis (PLS-DA) to distinguish between patient groups and identify differentially expressed variables. The correlation map of the first two components reveals a clear separation of the healthy individuals and group (solid and open grey circles respectively) from the pneumonia patients (**Fig. 5A**). The index values of the Variable Importance in Projection (VIP) from the PLS-DA were then used to identify 9 metabolites with VIP scores over one (**Fig. 5B**). However, the overall fit of this model was not robust (i.e. low Q^2^ values), indicating the quality of the fit varies a lot depending on the metabolite. Likewise, the R^2^ values were around 0.3 suggesting the components generated by the PLS regression did not summarize either the X or Y variables well. Thus, we revised this analysis by first centering and reducing the explanatory variables before starting the PLS-DA calculations (PLS-DA^VCR^). The quality of the PLS-DA^VCR^ was improved (i.e. Q² cumulative 0.083 to 0.378). While the Q² value is positive, thus has predictive relevance, it remains somewhat low suggesting the quality of the fit of this model varies a lot depending on the metabolite. The PLS-DA^VCR^ also improved the regression’s ability to summarize both the X and Y variables (i.e. R²Y 0.552 and R²X 0.483) resulting in better separation of pathogen groups (**Fig. 5C**). However, this produced a large number of metabolites, 35% of those identified, with VIP scores above 1 (**Fig. 5D**).

**Figure 5.**
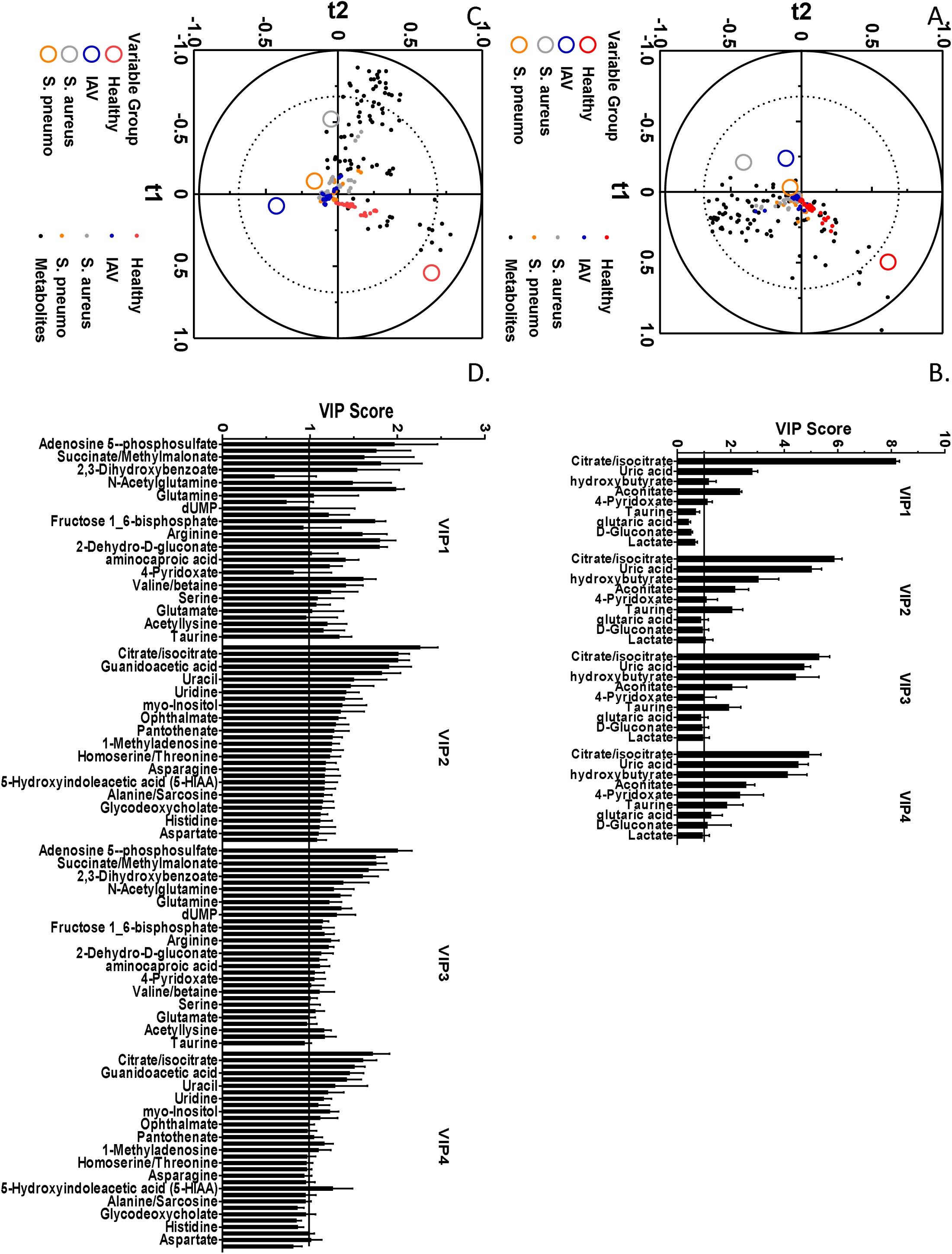
Identification of metabolites of import. Four component partial least squares discriminant analysis (PLS-DA) was used to identify metabolites with corresponding variable importance in the projection (VIP) values above 1. Correlation maps are depicted with variable groups and metabolite dispersion. Metabolites with the highest contribution to the model are graphed, with VIP score equal to one marked by a black line.

### Predictive Modeling

There were 291 variables including two demographics such as gender and age and 185 OTUs detected in urine samples, 17 cytokines, and 87 metabolites of 40 subjects. Note that we included OTUs that were observed for at least two subjects. First, we implemented multi-class classification with 5-folds cross validation to distinguish between four subject categories using a total of 291 predictors. However, none of the machine learning model provided desirable accuracy (all<47.5%).

We then implemented our recursive binary classification and variable selection approach using LASSO in three steps. To do that, we organized a separate dataset and an outcome variable for each step. For Model 1, we included all 40 subjects and the outcome variable were coded as 0 for ten heathy subjects and one for other 30. For Model 2, we included 30 subjects by excluding healthy ones and the outcome variable was coded as 1 for the 10 influenza subjects and 0 for the 10 *S. aureus* and 10 *S. pneumoniae* subjects. For Model 3, we included 20 subjects and outcome variable was coded as 0 for the 10 *S. aureus* and 1 for the 10 *S. pneumoniae* subjects. Initiallly, we compared Model 1 to the three traditional analysis methods above. We used four types of global analysis resulting in a total of 37 metabolites identified as potential biomarkers of pneumonia (**Table 4)**. Normalized peak intensities from each of these metabolites were then selected and analyzed individually using ANOVA followed by caparison of the means of healthy to pneumonia samples by Dunnett’s multiple comparisons test. Of the 37 metabolites identified with all the analysis only 18 metabolites were significantly altered by CAP (**Table 4 and S5A Fig.**). Using the individual manual analysis by Dunnetts, we then determined the rates of Type I and II errors from the grouped data analysis and the models **(Table 5**). None of the 6 metabolites identified by Tukey HSD failed the Dunnett’s test, but this method overlooked 12 metabolites (**Tables 4 and S5 Fig.**). Given the post hoc correction method was for false discovery rate, it is not surprising that this expression analysis resulted in no Type I error but high levels of Type II Errors (**Table 5**). The initial PLS-DA identified 9 metabolites with VIP > 1, of these only citrate and taurine showed significant differences (**Table 4 and S5A Fig.**). Further, the PLS-DA analysis produced the most, 16, Type II errors (**Table 5**). The PLS-DA^VCR^ analysis identified 31 metabolites with VIP > 1 (**Fig. 5D**). Nineteen of the metabolites identified with PLS-DA^VCR^ analysis passed the individual analysis thereby improving the Type II errors when compared to the PLS-DA. However, it misidentified 13 metabolites resulting in the largest number of Type I errors of any of the models (**Table 5**). Lasso Model 1 identified 7 potential biomarkers that distinguished healthy from CAP patients, all of which passed the Dunnett’s test (**Table 4 and S5 Fig.**). Model 1 also produced the least errors with more positive identifications (**Table 4**). Thus Lasso model 1 did not require reducing explanatory variables as was done in the PLS-DA^VCR^ analysis, that resulted in the greatest level of Type I errors, while producing the least Type I or II errors. It is important to note that in the first iteration of our model our model pulled out several predictions that are significantly altered and represent abundant metabolite markers in the urine.

**Table 4.** Comparisons of potential metabolite biomarkers identification methods.

| Metabolite | Tukey's<br>HSD | PLS-DA | PLS-DA<br>VR | Lasso<br>model | Dunnett's test |  |
| --- | --- | --- | --- | --- | --- | --- |
|  |  |  |  |  | Pass | Fail |
| 1-Methyladenosine |  |  | 1 |  | 1 |  |
| 2,3-Dihydroxybenzoate | 1 |  | 1 | 1 | 1 |  |
| 2-Dehydro-D-gluconate |  |  | 1 | 1 | 1 |  |
| 4-Pyridoxate |  | 1 | 1 |  |  | 1 |
| 5-Hydroxyindoleacetic acid (5-HIAA) |  |  | 1 |  | 1 |  |
| Acetyllysine |  |  | 1 |  | 1 |  |
| Aconitate |  | 1 |  |  |  | 1 |
| Adenosine 5--phosphosulfate | 1 |  | 1 | 1 | 1 |  |
| Alanine/Sarcosine |  |  | 1 |  |  | 1 |
| aminocaproic acid |  |  | 1 |  | 1 |  |
| Arginine |  |  | 1 |  | 1 |  |
| Asparagine |  |  | 1 |  |  | 1 |
| Aspartate |  |  | 1 |  |  | 1 |
| Citrate/isocitrate | 1 | 1 | 1 |  | 1 |  |
| D-Gluconate |  | 1 |  |  |  | 1 |
| dUMP |  |  | 1 | 1 | 1 |  |
| Fructose 1_6-bisphosphate |  |  | 1 |  | 1 |  |
| Glutamate |  |  | 1 |  |  | 1 |
| Glutamine |  |  | 1 |  |  | 1 |
| glutaric acid |  | 1 |  |  |  | 1 |
| Glycodeoxycholate |  |  | 1 |  |  | 1 |
| Guanidoacetic acid | 1 |  | 1 | 1 | 1 |  |
| Histidine |  |  | 1 |  |  | 1 |
| Homoserine/Threonine |  |  | 1 |  |  | 1 |
| hydroxybutyrate |  | 1 |  |  |  | 1 |
| Lactate |  | 1 |  |  |  | 1 |
| myo-Inositol |  |  | 1 |  |  | 1 |
| N-Acetylglutamine |  |  | 1 | 1 | 1 |  |
| Ophthalmate |  |  | 1 |  |  | 1 |
| Pantothenate |  |  | 1 |  | 1 |  |
| Serine |  |  | 1 |  |  | 1 |
| Succinate/Methylmalonate | 1 |  | 1 |  | 1 |  |
| Taurine |  | 1 | 1 | 1 | 1 |  |
| Uracil |  |  | 1 |  |  | 1 |
| Uric acid |  | 1 |  |  |  | 1 |
| Uridine | 1 |  | 1 |  | 1 |  |
| Valine/betaine |  |  | 1 |  | 1 |  |

**Table 5.** Errors per analysis method

|  | Error Type |  |
| --- | --- | --- |
|  | I | II |
| Tukey's HSD | 0 | 12 |
| PLS-DA | 7 | 16 |
| PLS-DA VCR | 13 | 0 |
| Model 1 | 0 | 11 |

We implemented LASSO logistic regression with fivefold cross-validation and a total of 13 OTUs, 2 cytokines, and 13 metabolites were found to be discriminating between different subject categories as listed in **Table 6**.

**Table 6.**
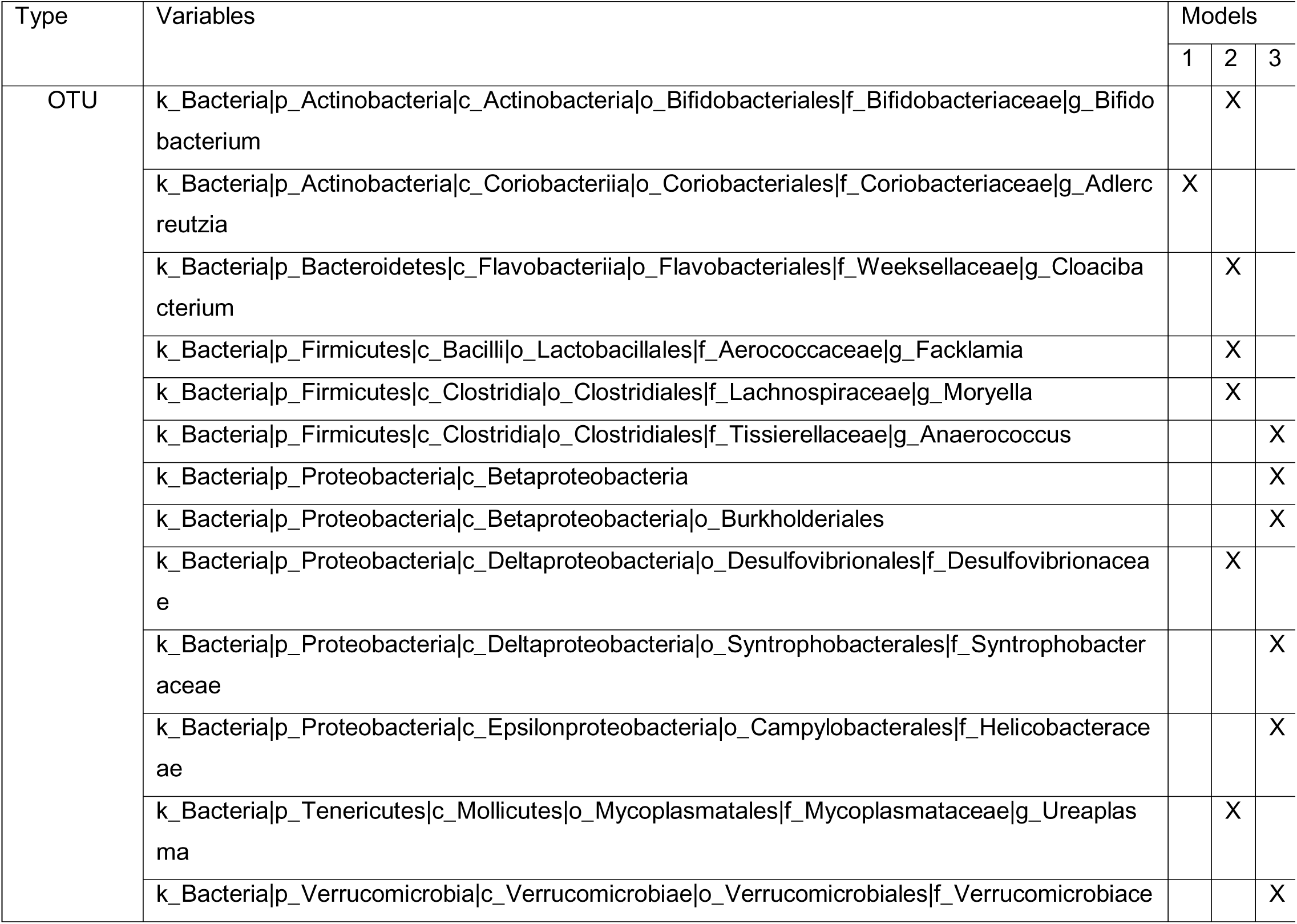
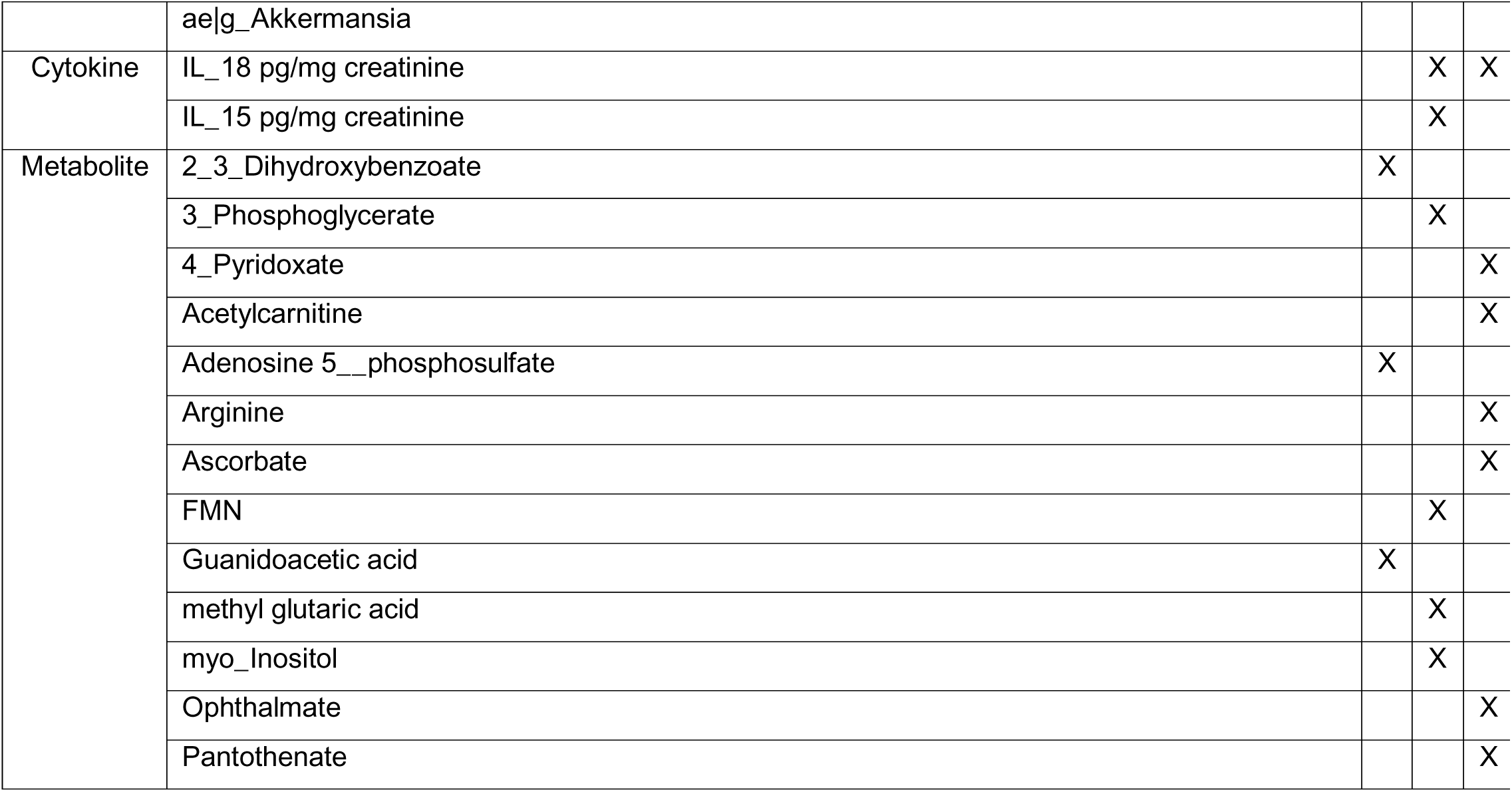
OTUs and Metabolites selected by LASSO

In Model, 1 we found one OTUs and three metabolites to distinguish heathy subjects from *S. aureus*, *S. pneumoniae*, and Influenza (Model 1 providing AUC with 95%CI of 0.98; 0.94-1.00), six OTUs, two cytokines, and four metabolites to distinguish influenza from *S. aureus* and *S. pneumoniae* (Model 2 providing AUC of 1.00), and six OTUs, one cytokine, and six metabolites to distinguish *S. aureus* from *S. pneumoniae* (Model 3 providing AUC of 1.00). The confusion matrices with performance indicators for each model is presented in **Table 7.**

**Table 7.**
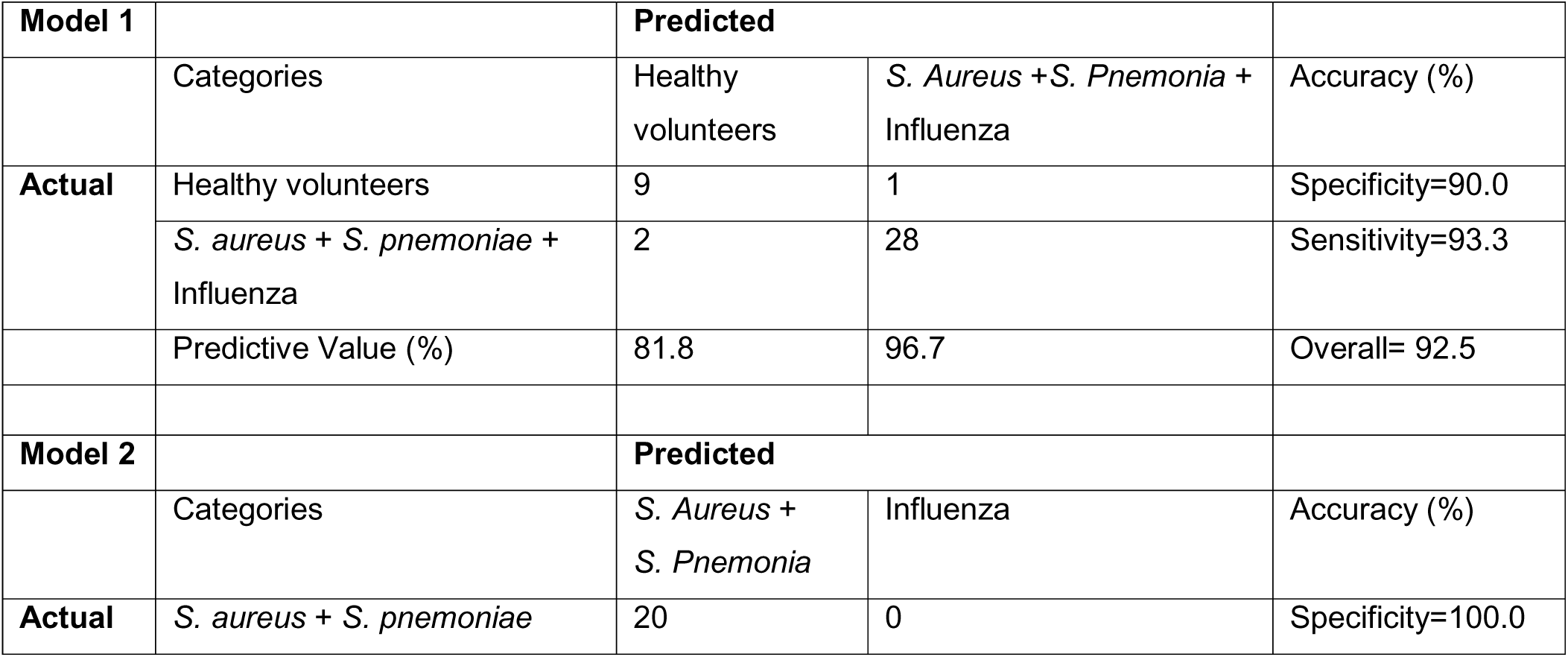
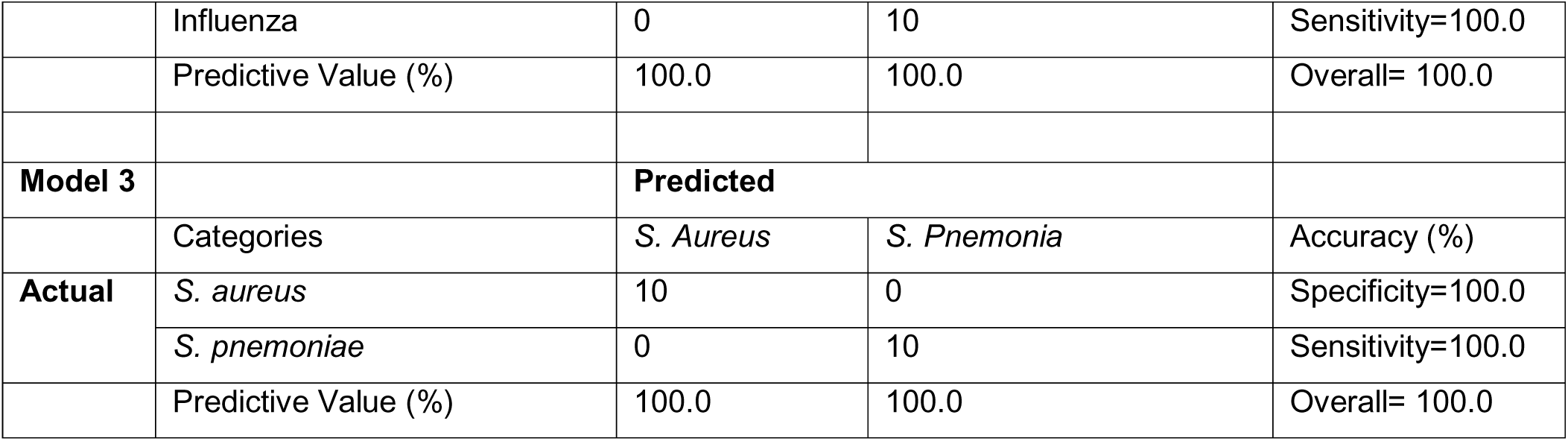
5-folds Cross-Validation LASSO Model Performances.

In recursive implementation of LASSO regression in three steps, we identified a total of 28 OTUs, cytokines, and metabolites to classify subjects into their actual categories. However, model 1 assumes the subject is not healthy and model 3 assumes the subjects not healthy nor influenza. Therefore, to develop a model that can be implemented on subjects without any assumption on their status, by using these selected 28 predictors, which is significantly smaller than the original dataset with 291 predictors, we readdressed the multi-class classification problem. We implemented various machine learning algorithms and found that Ensemble Method (Ensemble Method: Subspace, Learner Type: Discriminant, Number of Learners: 30, Subspace Dimension: 13) provided the highest overall classification accuracy of 85.0% **Table 8.**

**Table 8.** 5-folds Cross-Validation Ensemble Method Performance.

|  |  | **Predicted** |  |  |  |  |
| --- | --- | --- | --- | --- | --- | --- |
|  |  | Healthy volunteers | Influenza | *S. Pneumonia* | Healthy volunteers | Accuracy (%) |
| **Actual** | Healthy volunteers | 8 | 0 | 1 | 1 | 80.0 |
|  | Influenza | 0 | 9 | 1 | 0 | 90.0 |
|  | *S. pneumoniae* | 0 | 0 | 10 | 0 | 100.0 |
|  | *S. aureus* | 0 | 0 | 3 | 7 | 70.0 |
|  | Predictive Value (%) | 100.0 | 100.0 | 66.7 | 87.5 | Overall = 85.0 |

**Table 8** shows that most of the classification error is due to misclassification between two bacterial groups. When we merge two bacterial groups (**Table 9**), we found that one can distinguish between healthy, influenza and bacterial categories with very high accuracy of 92.5% and with perfect positive predictive values for healthy and influenza subjects.

**Table 9.** Performance of the final model when S. aureus and S. pneumoniae are merged into one group.

|  |  | Predicted |  |  | Accuracy (%) |
| --- | --- | --- | --- | --- | --- |
|  |  | Healthy volunteers | Influenza | Bacteria |  |
| **Actual** | Healthy volunteers | 8 | 0 | 2 | 80.0 |
|  | Influenza | 0 | 9 | 1 | 90.0 |
|  | Bacteria | 0 | 0 | 20 | 100.0 |
|  | Predictive Value (%) | 100.0 | 100.0 | 87.0 | Overall=92.5 |

## Discussion

The search for accurate predictors of infection and disease remains an important frontier in the era of omics and personalized medicine. In the setting of CAP, which is the leading cause of infectious related death in the United States, various methods have been employed using serum and other clinical samples to confirm and determine severity of respiratory infection, including the quantification of cytokines and eicosanoids. However, these approaches have achieved limited sucess and are not widely implemented for patient care. For the clinician, even the basic differential diagnosis between viral vs bacterial pathogens remains difficult in CAP cases due to the overlapping symptomatic presentations. Considering the dynamic nexus of host immunological, metabolic, and microbial networks, we moved beyond the search for single or limited numbers of biomarkers by instead comprehensively profiling urine cytokines, the microbiome, and the metabolome in samples collected from newly admitted pneumonia patients with either influenza or bacterial pathogens compared with healthy volunteers. Urine was chosen as a non-invasive sample since it is readily obtainable in the in-patient and out-patient setting.

There have been few studies to determine the utility of measuring cytokine markers in urine as potential biomarkers of infectious disease states. Out of the 34 cytokines we measured in urine, 11 were significantly different between groups. Patients with bacterial pneumonia exhibited the greatest elevation and number of cytokines in urine that differed from healthy controls; *S. aureus* pneumonia patients differed in all 11 cytokines with IL-4, IL-15, Gro-α, MIP-1α and SDF-1 uniquely elevated in those patients. Whereas patients with influenza exhibited the lowest levels of cytokines detectable in urine that differed compared to healthy controls. In several studies IL-6 levels in the serum have been found to correlate with either bacterial infections or disease severity in patients with CAP. ^44–47^ Our results also detected increased IL-6 in the urine which was elevated in all pneumonias compared to healthy controls. However, based on the predictive modeling, IL-15 and IL-18 in combination with the metabolome and microbiome data may be more useful for distinguishing bacterial vs viral pneumonias.

Analysis of the urine microbiome demonstrated a complex community under both healthy and infectious states, including the presence of slow growing *Lactobacillus* and *Corynebacterium*, consistent with recent reports on urine microbiome. While urinary tract bacterial populations were historically overlooked outside of the context of infection, next generation sequencing techniques enabled culture independent insights into these communities. Recent findings demonstrate major shifts in the urine microbiome community under diseases of the urinary tract, including ^48^ urolithiasis and certain cancers, however investigation of the urine microbiome as signatures of disease at distant body sites has not been employed. We observed elevated diversity and evenness in urine samples from pneumonia patients compared with healthy volunteers, including consistently elevated *Enterobacteriaceae* and *Clostridium*. Furthermore, community composition data suggested greater dissimilarity in the urine microbiome in patients with bacterial pneumonia than in those with influenza (**Fig. 2A-C**).

Similar to the microbiome, the metabalome of CAP patients clustered from healthy volunteers (**Fig. 3 and Fig. 4**), suggesting a divergence in metabolite profiles under an infectious state. Drivers of this differential clustering included loss of numerous metabolites, including citrate/isocitrate, succinate, guanidoacetic acid, N-acetylglutamine, among others, compared with healthy volunteers. No metabolites were specific to influenza infection alone compared with the other groups. However, methyladenosine, uridine and 2-dehydro-D-gluconate were elevated under bacterial infection compared with influenza and healthy controls. These divergent profiles could be useful in determining pathogen kingdom.

Since the microbiome and metabolite profiles are influenced by the environment, nutrition, age, and lifestyle of the host, in addition to genetics, these concatenated profiles provide a unique snapshot of individual health. ^19, 22, 31, 23–30^ Indeed, while these complex profiles can be examined independently, changes in the collective abundance patterns of metabolites and microbes may indicate deeper homeostatic disturbances, which may be reflected through changes in interleukin signaling. The membership of the bodies microbial communities have dynamic interconnected relationships with one another and the host that change under states of disease and stress. Therefore, the microbiome and metabolome complement to serve as personalized readout of individual health. The ability to detect rapid measurable changes in these profiles in response to challenges, such as infection, would be a novel systems biology approach to personalized medicine. For instance, the components of the metabolome and microbiome are physically or stoichiometrically co-related, leaving precise abundance patterns that may accurately reflect discreet pathophysiological states ^23, 34^. Utilization of meta-biomarkers, such as the microbiome and metabolome, would represent a distinct shift away from the majority of clinical biomarkers currently in use, even in the era of genomics, transcriptomics, and proteomics ^23, 34^.

After combining all urine meta-biomarkers, totaling 385 data points, we performed machine learning models by implementing LASSO logistic regression in 3 unique models. Model 1 aimed to distinguish healthy subjects from pneumonia; Model 2 to distinguish between bacterial (*S aureus* and *S pneumoniae*) pathogens from Influenza; and Model 3 to distinguish between *S aureus* and *S pneumoniae*. For each predictive model, we implemented a 5-fold cross validation process to avoid overfitting. Specifically, the data were split into five distinct folds where 4 folds were used for model testing and the remaining for validation. By repeating this process five times by changing the test fold, we identified a total of 28 predictors, including two cytokines, 13 microbial taxa, and 13 metabolites that provided a predictive power of 92.5% in distinguishing patient groups.

There are several limitations to our study, including the total sample size of 40 individuals. While we were able to detect consistent changes in our meta-biomarkers, larger studies with greater numbers and groups that included other pathogens may improve the resolution of our predictive models in determining unique signatures of pneumonia or other respiratory diseases. Another limitation was related to a characteristic of the clinical standard of care, where all pneumonia patients in this study were placed on antibiotics upon admission to the hospital. Future studies may attempt to compare urine samples collected from individual before and after the implementation of antibiotics. On the other hand, the inclusion of patients with influenza acted as a unique control group for the bacterial groups, since all patients were placed on antibiotics. We observed large perturbations in the meta-biomarkers in bacterial groups compared with the influenza group, suggesting that the changes were indeed driven by the pathogen and not a general response to infection.

Here we describe a comprehensive profile of urine meta-biomarkers, including the microbiome, metabolome, and cytokines in pneumonia patients. Using these biomarkers, we achieved high success in predicting pneumonia pathogens. Depending on the infectious pathogen identified in each patient, distinctly different immune profiles were observed in cytokine profiles, and even larger shifts were observed in the metabolite and microbiome profile, especially in response to bacterial infections. This study provides a proof of concept that urine samples, which are easily accessible in outpatient and inpatient settings, could provide additional diagnostic insights to patient infectious status and future risk factor for complication.

## Materials and Methods

### Ethics Statement

The usage of human samples was approved and performed in accordance with the regulations and guidelines set by the Univeristy of Louisville Insitutional Review Boards and the Human Subjects Protection Program. Samples were obtained from the University of Louisville Infectious Disease Biorepository (IRB # 13.0001) and de-identified metadata were used for analysis under the Biomarkers study (IRB # 17.0601). All patients provided written informed consent for sample biorepository storage and subsequent use in research studies.

### Sample Processing

Urine samples were collected using sterile technique and were aliquoted separately for cytokines, microbiome analysis, and metabolites^15, 16, 36^. For multiplex cytokine assays and microbiome analysis, the urine samples were centrifuged at 10,000xg for 10 min and then analyzed as described below. For Metabolite analysis, samples (50 ul for urine) were extracted with 1.3 mL of extraction solvent (40:40:20 HPLC grade methanol, acetonitrile, water with 0.1% formic acid) pre-chilled to 4°C in a cold room and incubated for 20 min at −20°C. The samples were centrifuged for 5 min (16.1 rcf) at 4°C. The supernatants were transferred to new 1.5 mL centrifuge tubes and pellets were resuspended with 200 µL of extraction solvent. Extraction was allowed to proceed for 20 min at −20°C and all supernatants collected in glass vials. Vials containing the collected supernatant were dried under a stream of N2 until all the extraction solvent had been evaporated. Residue was resuspended in 300 µL of sterile water and transferred to 300 µL autosampler vials. Samples were immediately placed in autosampler trays for mass spectrometric analysis.

### Multiplex for cytokines and statistical analysis

The levels of a panel of inflammatory mediators in urine samples were measured using a 34-plex ProcartaPlex™ Multiplex Immunoassay according to manufacturer’s instructions (Invitrogen, Carlsbad CA, USA). Cytokine standards were prepared to determine the concentration of cytokines in the samples. The samples were run on a Millipore Magpix instrument and analyzed with xPONENT 4.2 software. For data analysis, a five-parameter logistic curve fitting method was applied to the standards and the sample concentrations extrapolated from the standard curve. The results were normalized to creatinine as measured by Creatinine Detection Kit (Thermofisher, Waltham, MA).

Kruskal-Wallis test was used to compare median level of cytokine among the four groups of samples. The median cytokine levels between urine samples from health volunteers and those from influenza, *S. pneumo* or *S. aureus* were tested via Wilcoxon sum rank test. The p values were not adjusted for multiple comparisons. All analyses were performed using R-3.4.0 (https://www.R-project.org/)

### Microbial DNA isolation

Human urine samples (100uL) were centrifuged at 10,000 x g for 10 minutes, supernatant was carefully removed, and 500 uL of extraction buffer (50mM Tris (pH 7.4), 100mM EDTA (pH 8.0), 400mM NaCl, 0.5% SDS) containing 20uL proteinase K (20 mg/ml, Cat# 03115887001, Roche) was added to each tube^15, 49^. 0.1-mm-diameter zirconia/silica beads (BioSpec Products, Bartlesville, OK, USA) were added to the extraction tubes and a Mini-Beadbeater-8 cell disrupter (BioSpec Products) for 2 x 1 minute to lyse cells. After overnight incubation at 55°C with agitation, extraction with phenol:chloroform:isoamyl alcohol, and precipitation with ethanol were performed. Isolated DNA was dissolved in nuclease-free water and stored at −80°C.

### 16S rRNA-based PCR, ilumina library preparation, and data analysis

To assess total 16S copy numbers, 2uL of isolated DNA was used in quantitative PCR analysis using 16S primers (Forward: 5’-TCCTACGGGAGGCAGCAGT-3’; Reverse: 5’-GGACTACCAGGGTATCTAATCCTGTT-3’) and an in-house standard to generate a standard curve. To assess bacterial community structure, primers specific for 16S rRNA V4-V5 region (Forward: 338F: 5’-GTGCCAGCMGCCGCGGTAA-3’ and Reverse: 806R: 5’-GGACTACHVGGGTWTCTAAT-3’) that contained Illumina 3’ adapter sequences, as well as a 12-bp barcode, were used. Sequences were generated by an Illumina MiSeq DNA platform at Argonne National Laboratory and analyzed by the program Quantitative Insights Into Microbial Ecology (QIIME). ^50^Operational Taxonomic Units (OTUs) were picked at 97% sequence identity using open reference OTU picking against the GreenGenes database. OTUs were quality filtered based on default parameters set in the open-reference OTU command in QIIME and sequences were rarified to an equal sampling depth of 8,000 reads per sample. Representative sequences were aligned via PyNAST, taxonomy assigned using the RDP Classifier, and a phylogenetic tree was built using FastTree. Processed data were then imported into Calypso 8.84 for further analysis and data visualization. ^51^ Calypso data underwent total sum normalization with Hellinger square root transformation. Alpha diversity was assessed using observed Shannon index and Eveness. Network analyses were generated with Speaman’s correlations. Positive correlations were FDR-adjusted at p < 0.05 aand presented as network edges. OTUs generated in QIIME were finally analyzed using linear discriminant analysis (LDA) effect size (LEfSe) where non-parametric factorial Kruskal-Wallis sum-rank testing (*p* < 0.05) identified significantly abundant taxa followed by unpaired Wilcoxon rank-sum test to determine LDA scores >2 was considered significant.

### Metabolite analysis

#### UPLC–HRMS metabolomics analysis

Samples placed in an autosampler tray were kept at 4 °C. A 10 µL aliquot was injected through a Synergi 2.5 micron reverse-phase Hydro-RP 100, 100 x 2.00 mm LC column (Phenomenex, Torrance, CA) kept at 25 °C. The eluent was introduced into the MS via an electrospray ionization source conjoined to an Exactive™ Plus Orbitrap Mass Spectrometer (Thermo Scientific, Waltham, MA) through a 0.1 mm internal diameter fused silica capillary tube. The mass spectrometer was run in full scan mode with negative ionization mode with a window from 85 – 1000 m/z. with a method adapted from Lu et al., 2010 ^52^. The samples were run with a spray voltage was 3 kV. The nitrogen sheath gas was set to a flow rate of 10 psi with a capillary temperature of 320°C. AGC (acquisition gain control) target was set to 3e6. The samples were analyzed with a resolution of 140,000 and a scan window of 85 to 800 m/z for from 0 to 9 minutes and 110 to 1000 m/z from 9 to 25 minutes. Solvent A consisted of 97:3 water:methanol, 10 mM tributylamine, and 15 mM acetic acid. Solvent B was methanol. The gradient from 0 to 5 minutes is 0% B, from 5 to 13 minutes is 20% B, from 13 to 15.5 minutes is 55% B, from 15.5 to 19 minutes is 95% B, and from 19 to 25 minutes is 0% B with a flow rate of 200 µL/min ^52^.

Files generated by Xcalibur (RAW) were converted to the open-source mzML format ^53^ via the open-source msconvert software as part of the ProteoWizard package ^54^. Maven (mzroll) software, Princeton University ^55, 56^ was used to automatically correct the total ion chromatograms based on the retention times for each sample. Metabolites were manually identified and integrated using known masses (± 5 ppm mass tolerance) and retention times (Δ ≤ 1.5 min). Unknown peaks were automatically selected via Maven’s automated peak detection algorithms.

Multivariate statistical analysis for the MS/MS data was performed using XLSTAT software (Addinsoft, New York, NY) interfaced with Excel (Microsoft Corporation, Redmond, WA). The average coefficient of variation (C.V.) was 0.395 (+/- 0.211). Thus an inclusion criterion for technical replicates were applied based on C.V. ≤ 0.5 resulting in 11 exclusion (i.e. 11 technical replicates in duplicate and 29 in triplicate) resulting in C.V. average of 0.288 (+/- 0.114). To ensure that observations were directly comparable and to account for the biofluid concentration peak intensity was normalized to creatinine (these data were compared to unnormalized data to make sure there was no masking of biologically relevant changes by normalization). To evaluate the group trends, sample uniformity and identify potential outliers unsupervised multivariant principal component analysis (PCA) was employed. The variation were explained by F1 and F2 with a cumulative percent variability of 78.558 and a marginal increase to 85.958% with the addition of F3. These data were then independently k-means clustered followed by ascendant hierarchical clustering based on Euclidian distances. The data matrix’s was rearranged according to the corresponding clustering with similarity proportional to a closer spatial relationship for patient sample columns and metabolite rows. 21 metabolites with less than 0.25 standard deviation were eliminated to simplify the graph. These clusters were also represented via a dendrogram displayed vertically for metabolites and another horizontally for patients. The data values of the permuted matrix were replaced by corresponding color intensities based on interquartile range with color scale of red to green through black resulting in a heat map. Patient identifiers and risk categories were replaced by color bars. XLSTAT expression analysis was then used to determine metabolite significance between groups using one-way ANOVA with Benjamini-Hochberg post hoc correction and found to have with significant differences using Tukey’s honest significance test (Tukey HSD) for multiple comparisons. Partial least squares discriminant analysis (PLS-DA) was then applied to separate patient groups and identify metabolites with corresponding variable importance in the projection (VIP) values above 1. The 4 component PLS-DA was then rerun with the variables centered and reduced prior to analysis to improve the model quality and identify the corresponding VIP. Each identified metabolite was then analyzed individually using ANOVA then the means of pneumonia samples were then compared to healthy controls (Dunnett’s multiple comparisons test) prior to and post outlier removal performed with PRISM software (Graphpad, San Diego, CA).

### Predictive Modeling

We implement predictive modeling approach to distinguish between four subject categories (healthy, *S.Aureus, S. Pneumoniae*, and Influenza) using identified operational taxonomic units (OTUs) and metabolites from urine samples. First, all 40 subjects were analyzed to identify OTUs. Sample decompositions were normalized in a way that sum of all detected OTUs are equal to 1 for each subject. We then combined identified OTUs, cytokines and metabolites as predictors of four subject categories. Our first approach is to implement multi-class classification using various machine learning algorithms such as random forest, ensemble trees, support vector machines, k-nearest neighborhood. However, small sample size (n=40), multiple output categories (4 subject groups) and expected larger number of predictors (OTUs and metabolites) made predictive modeling very challenging. Therefore, as an alternative approach, we implemented recursive binary classification in three steps and obtained three different models at each step. First, Model 1 is to distinguish healthy subjects from the other three disease categories, Model 2 to distinguish between bacterial *(S. Aureus* and *S. Pneumoniae*) disease from Influenza, and Model 3 to distinguish between S. Aureus and *S. Pneumoniae.* Considering the small sample size and large number of predictors, for each of the three models, we first implement Least Absolute Shrinkage and Selection Operator (LASSO) ^57^ logistic regression ^58^. LASSO is statistical method retraining strong features of both subset selection and ridge regression. It implements ordinary least squares subject to sum of absolute values of the regression coefficients being less than a predetermined constant value. Logistic regression LASSO is an extension of LASSO for an output variable with binomial distribution.

By implementing LASSO, some of the regression coefficients are shrink to take a valued of zero therefore only variables with non-zero regression coefficients remain in the model. By taking advantage of LASSO, we will first identify OTUs and metabolites that are the most effective in distinguishing between our subject categories in Model 1, Model 2, and Model 3. Next, we combined all selected OTUs and metabolites in readdress multi-class classification problem using the machine learning approaches mentioned.

For each predictive model, we will implement 5-folds cross validation process to avoid overfitting. That is, the entire that is split into five distinct folds and a model built on using four folds of data and tested on the remaining fold. By repeating this process five times by changing the test fold, we obtain a predict class labels for each subject using a model that is trained on other subjects. We will compare different machine learning algorithms based on model performance statistics such as specificity, sensitivity, and positive and negative predictive value.

## Acknowledgement

Metabolomic extraction and mass spectrometric analyses were performed at the Biological and Small Molecule Mass Spectrometry Core, University of Tennessee, Knoxville, TN with the assistance of Dr. Shawn R. Campagna, Dr. Hector F. Castro, Sara Howard and Eric Tague. We thank members of the ISHPS Pneumonia Working Group, Reba Umberger, Amber Smith and Robert Williams for their input into experimental design and data analysis.

---

**Supplemental Figure 1.**
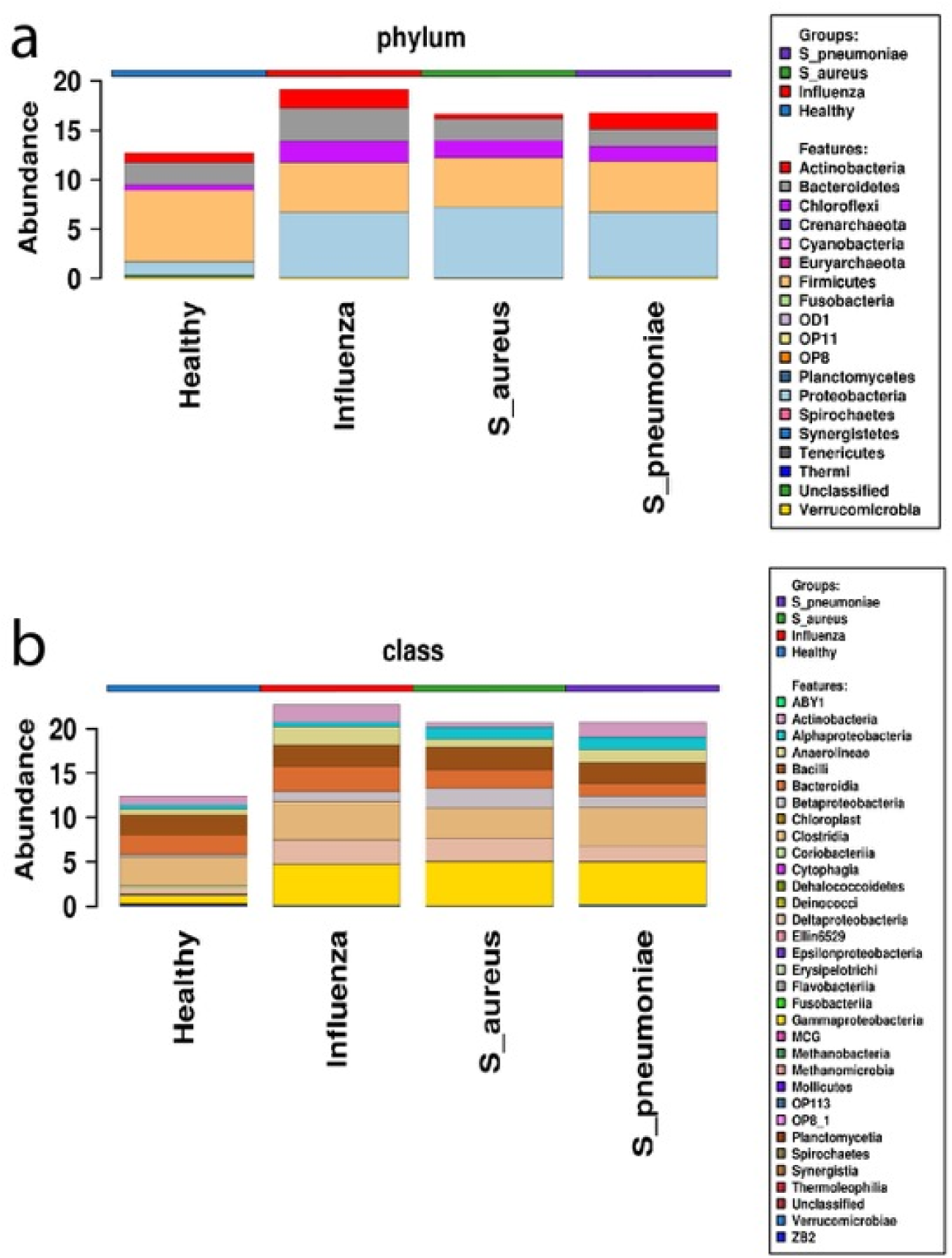
Microbiome taxonomic composition by group shown at the (a) phylum and (b) class level.

**Supplemental Figure 2.**
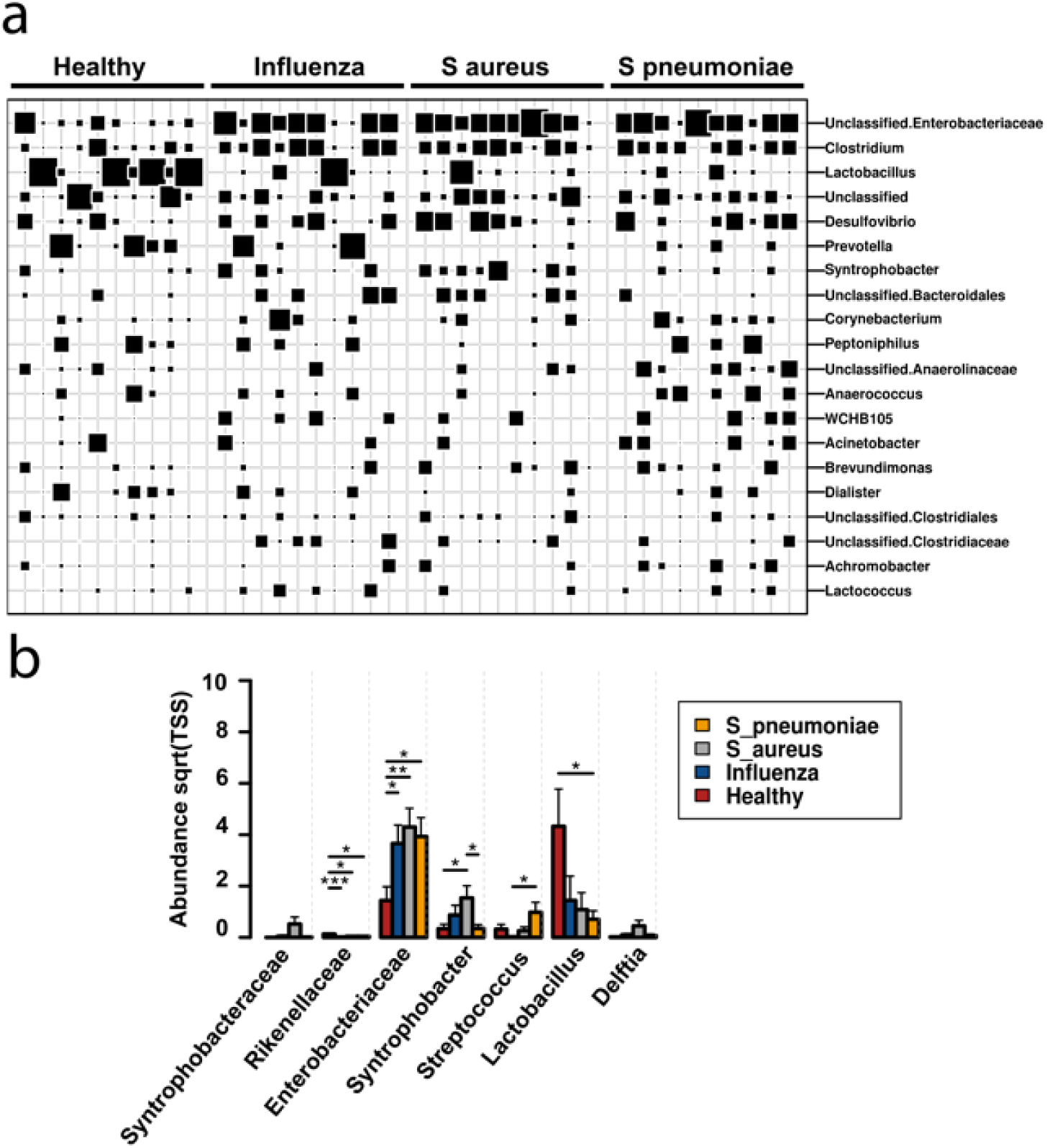
(a) Box plot shows relative abundance of taxa across samples within each experimental group. (b) Anova analysis of significantly altered taxa at the family level. * P < 0.05; ** P < 0.01; *** P < 0.005.

**Supplemental Figure 3.**
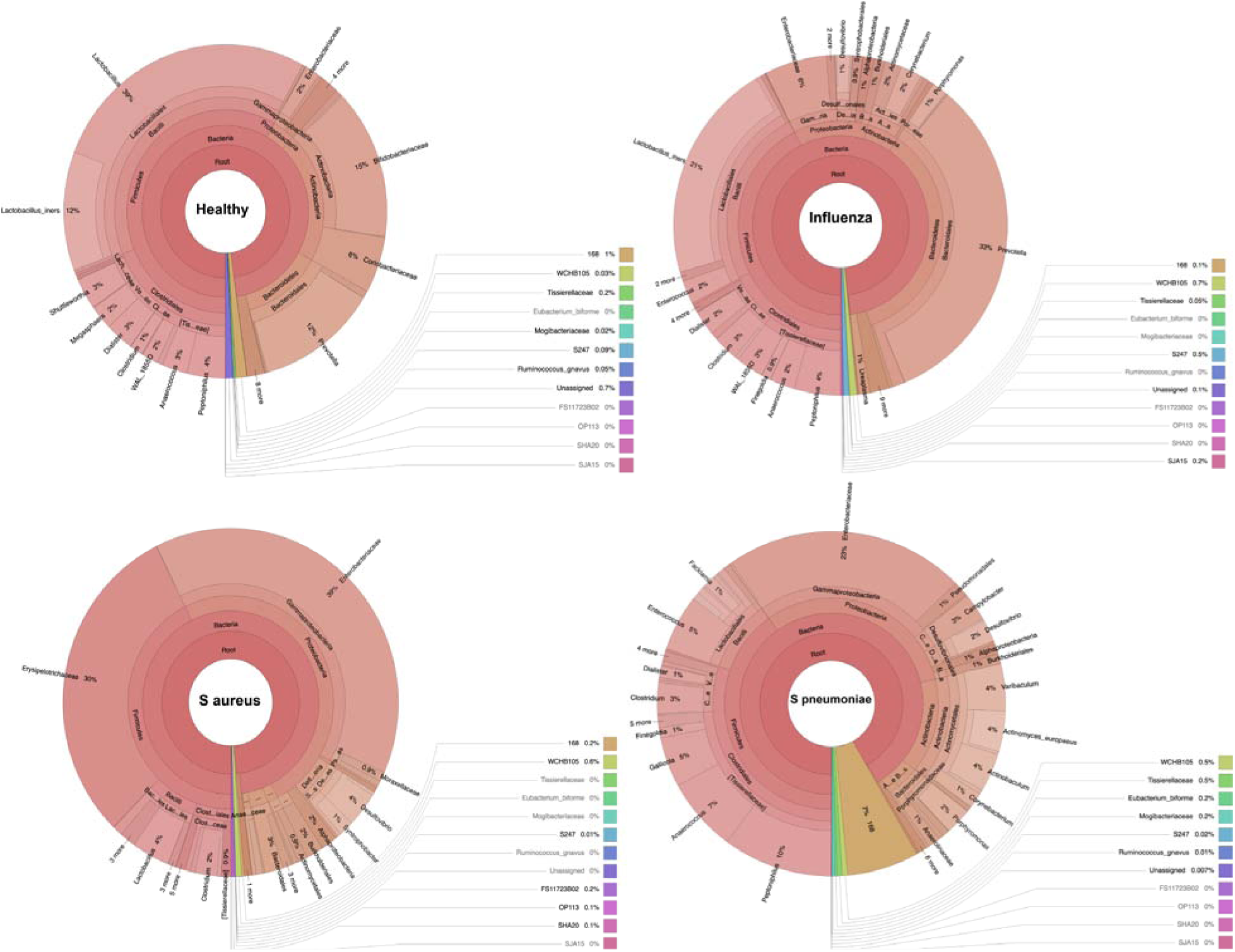
Krona plots showing the community composition at each taxonomic hierarchy for each experimental group.

**Supplemental Figure 4.**
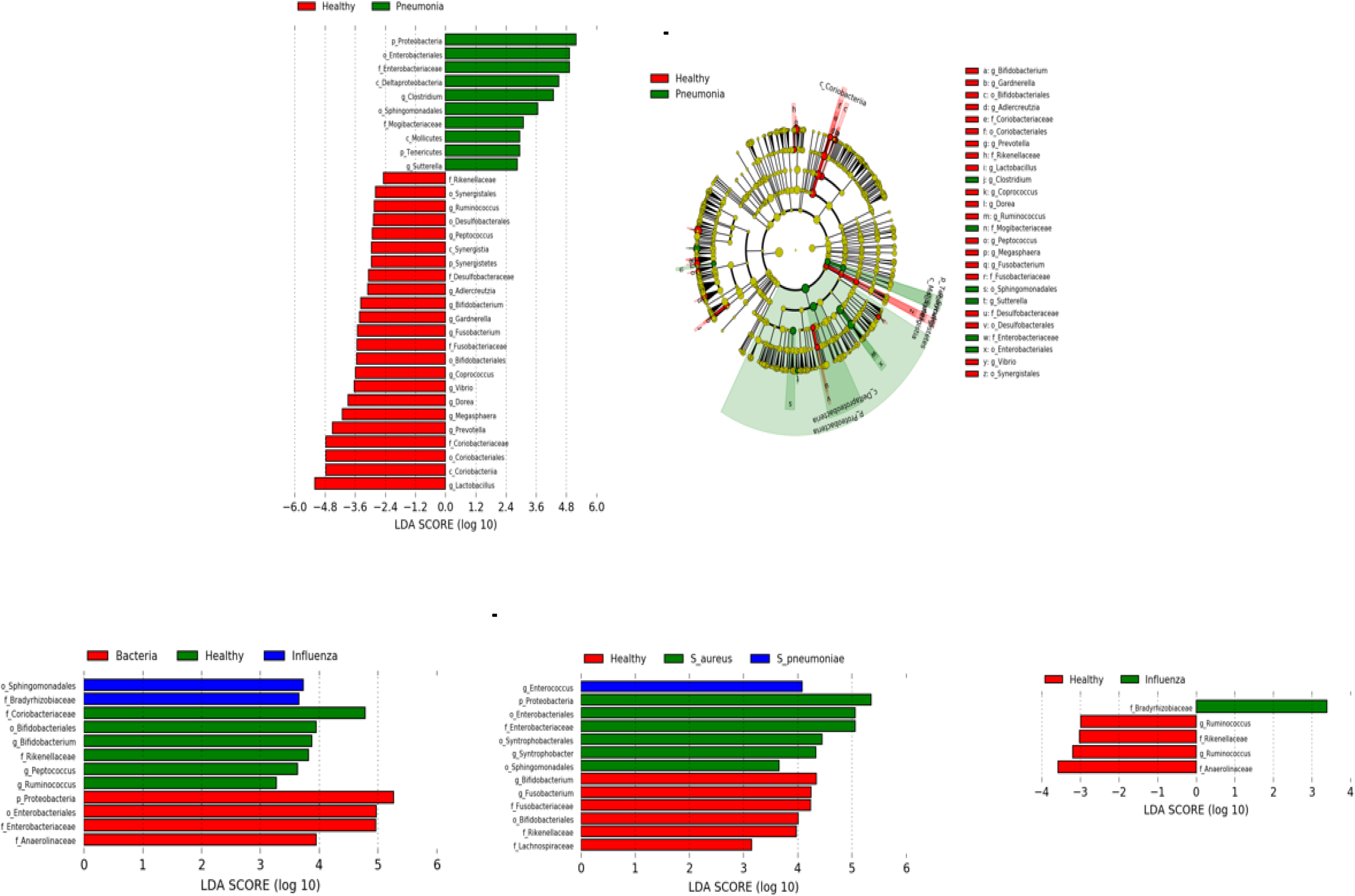
(a) Linear Discriminant Analysis of Effect Size (LefSe) of taxa enriched in healthy vs all pneumonia patients. (b) Dendrogram displaying the significantly altered taxa shown in panel a between healthy and pneumonia patients. (c) LefSe of taxa enriched in Healthy vs bacteria vs influenza patients. (d) LefSe of taxa enriched in healthy vs S aureus vs S pneumonia patients. (e) LefSe of taxa enriched in healthy vs influenza patients.

**Sup Fig 5.**
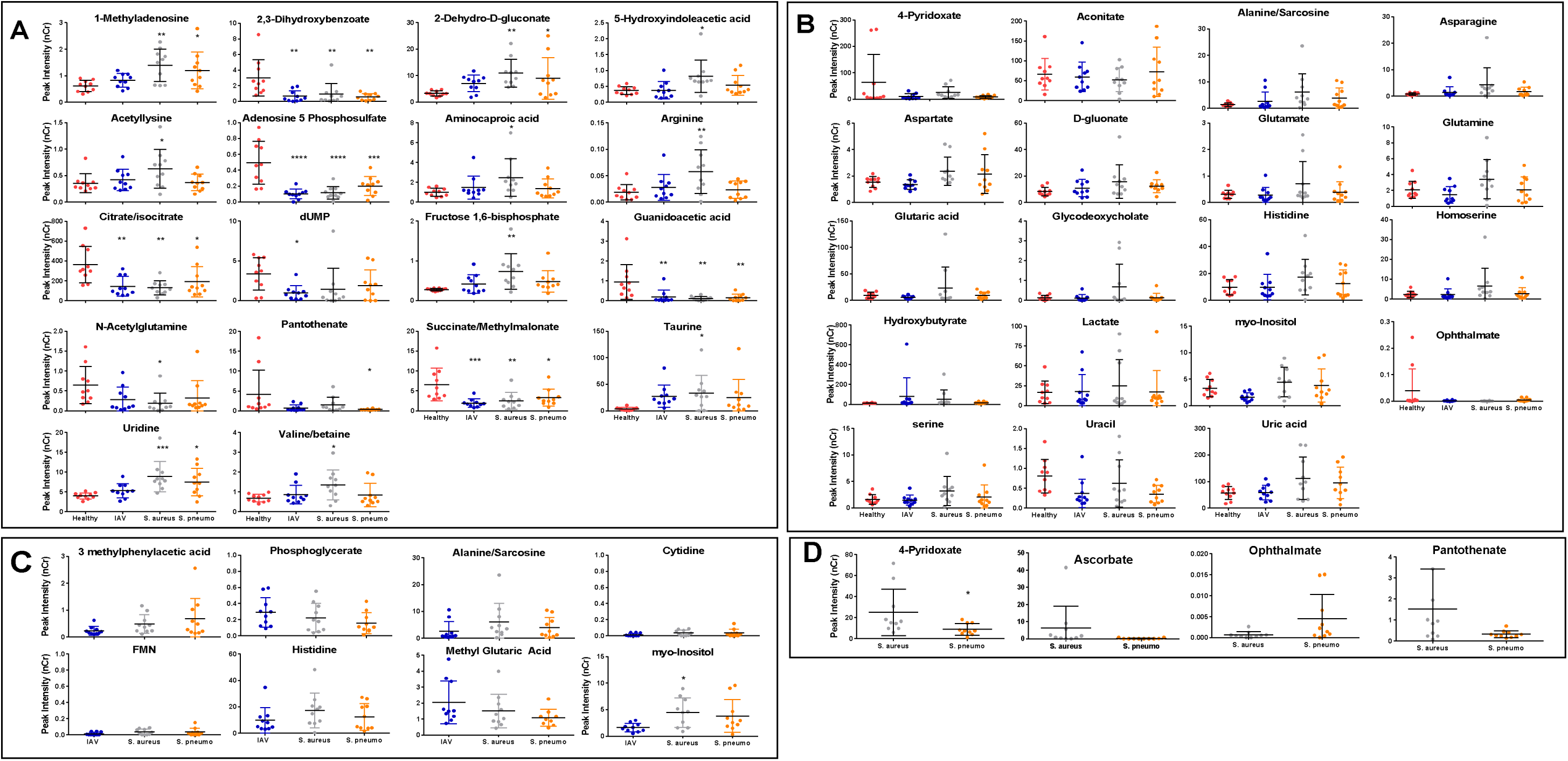
Individual metabolite analysis. Samples were subjected to analysis by group (means) using one-way ANOVA with Benjamini-Hochberg post hoc correction and considered for further testing if found to have with significant differences using Tukey’s honest significance test (Tukey HSD) for multiple comparisons. Each potential metabolite was then analyzed individually using ANOVA and the means compared to healthy controls using one-way ANOVA with Dunnett’s multiple comparison test (A & B). Infected patients were compared using one-way ANOVA with Tukey’s multiple comparisons test (C). Bacterial infections were compared using unpaired T test (D). Asterisks indicate significance as follows: p-value < 0.05 (*), p-value < 0.01(**), p-value < 0.001 (***), and p-value < 0.0001 (****).

